# MCU-enriched dendritic mitochondria regulate plasticity in distinct hippocampal circuits

**DOI:** 10.1101/2023.11.10.566606

**Authors:** Katy E. Pannoni, Quentin S. Fischer, Renesa Tarannum, Mikel L. Cawley, Mayd M. Alsalman, Nicole Acosta, Chisom Ezigbo, Daniela V. Gil, Logan A. Campbell, Shannon Farris

**Affiliations:** Fralin Biomedical Research Institute at Virginia Tech Carilion, Center for Neurobiology Research, Roanoke, Virginia; Graduate Program in Translational Biology, Medicine, and Health, Virginia Tech, Blacksburg, Virginia; Department of Biomedical Sciences & Pathobiology, Virginia-Maryland College of Veterinary Medicine, Virginia Tech, Blacksburg, Virginia; Virginia Tech Carilion School of Medicine, Roanoke, Virginia

**Keywords:** Mitochondrial calcium uniporter, Hippocampal CA2, Synaptic plasticity, Mitochondrial morphology, Dendrites, Spines

## Abstract

Mitochondria are dynamic organelles that are morphologically and functionally diverse across cell types and subcellular compartments in order to meet unique energy demands. Mitochondrial dysfunction has been implicated in a wide variety of neurological disorders, including psychiatric disorders like schizophrenia and bipolar disorder. Despite it being well known that mitochondria are essential for synaptic transmission and synaptic plasticity, the mechanisms regulating mitochondria in support of normal synapse function are incompletely understood. The mitochondrial calcium uniporter (MCU) regulates calcium entry into the mitochondria, which in turn regulates the bioenergetics and distribution of mitochondria to active synapses. Evidence suggests that calcium influx via MCU couples neuronal activity to mitochondrial metabolism and ATP production, which would allow neurons to rapidly adapt to changing energy demands. Intriguingly, MCU is uniquely enriched in hippocampal CA2 distal dendrites relative to neighboring hippocampal CA1 or CA3 distal dendrites, however, the functional significance of this enrichment is not clear. Synapses from the entorhinal cortex layer II (ECII) onto CA2 distal dendrites readily express long term potentiation (LTP), unlike the LTP- resistant synapses from CA3 onto CA2 proximal dendrites, but the mechanisms underlying these different plasticity profiles are unknown. We hypothesized that enrichment of MCU near ECII-CA2 synapses promotes LTP in an otherwise plasticity-restricted cell type. Using a CA2-specific MCU knockout (cKO) mouse, we found that MCU is required for LTP at distal dendrite synapses but does not affect the lack of LTP at proximal dendrite synapses. Loss of LTP at ECII-CA2 synapses correlated with a trend for decreased spine density in CA2 distal dendrites of cKO mice compared to control (CTL) mice, which was predominantly seen in immature spines. Moreover, mitochondria were significantly smaller and more numerous across all dendritic layers of CA2 in cKO mice compared to CTL mice, suggesting an overall increase in mitochondrial fragmentation. Fragmented mitochondria might have functional changes, such as altered ATP production, that might explain a deficit in synaptic plasticity. Collectively, our data reveal that MCU regulates layer-specific forms of plasticity in CA2 dendrites, potentially by maintaining proper mitochondria morphology and distribution within dendrites. Differences in MCU expression across different cell types and circuits might be a general mechanism to tune the sensitivity of mitochondria to cytoplasmic calcium levels to power synaptic plasticity.

**MAIN TAKE HOME POINTS:**

- The mitochondrial calcium uniporter (MCU) regulates plasticity selectively at synapses in CA2 distal dendrites.
- The MCU-cKO induced LTP deficit correlates with a trending reduction in spine density in CA2 distal dendrites.
- Loss of MCU in CA2 results in ultrastructural changes in dendritic mitochondria that suggest an increase in mitochondrial fragmentation. These ultrastructural changes could result in functional consequences, such as decreased ATP production, that could underlie the plasticity deficit.
- Dendritic mitochondrial fragmentation in MCU cKO occurred throughout the dendritic laminae, suggesting that MCU is dispensable for establishing layer-specific mitochondrial structural diversity.

## INTRODUCTION

Mitochondria dynamically regulate many critical cellular functions, including energy production and calcium buffering, to meet the unique demands of different cell types (Pekkurnaz & Wang, 2022; Sprenger & Langer, 2019; Fecher et al., 2019). Even within a cell, mitochondria display remarkable heterogeneity across subcellular compartments (Faitg et al., 2021; Lewis et al., 2018; Pannoni et al., 2023), which is especially critical for highly polarized cells such as neurons. The extent to which mitochondrial diversity influences cell-specific functions remains an open question. Mitochondrial dysfunction has been described in many neurological disorders, including Alzheimer’s disease (Ashleigh et al., 2023; Naia et al., 2023; Mary et al., 2023), schizophrenia (Ben-Shachar, 2017; Prabakaran et al., 2004; Ene et al., 2023), autism spectrum disorder (Rossignol and Frye, 2012), depression (Bansal et al., 2016) and bipolar disorder (Kato et al., 2000). For this reason, it is critical to understand the relationship between mitochondria and neuronal function. In neurons, there is growing evidence demonstrating the importance of mitochondria for synapses and plasticity (Devine and Kittler 2018; Tang and Zucker 1997; Li et al., 2004; Smith et al., 2016; Stowers et al., 2002). In particular, dendritic mitochondria in cultured hippocampal neurons are critical for maintaining spine density (Li et al 2004) and for powering local-translation-dependent structural plasticity (Rangaraju et al., 2019). In acute slices, disrupting mitochondrial fission impairs long-term potentiation (LTP) at CA3-CA1 Schaffer collateral synapses (Divakaruni et al., 2018). However, the molecular mechanisms regulating the cross talk between mitochondria and synapses are not well understood.

The mitochondrial calcium uniporter (MCU) is a channel that allows calcium flux across the inner mitochondrial membrane into the matrix of the mitochondria (Baughman et al., 2011, Stefani et al., 2011; Chaudhuri et al., 2013), where it has potentially far-reaching effects on mitochondrial form and function (Llorente-Folch et al., 2015; Zhao et al., 2015; Rizzuto et al., 2012; Wescott et al., 2019). While this has been extensively studied in pathological conditions and in culture (Qiu et al., 2013; Llorente-Folch et al., 2015; Zhao et al., 2015; Liang et al., 2014), little is known about the role mitochondrial calcium uptake plays in intact neural circuits. It’s thought that calcium influx via MCU may be a mechanism to couple neuronal activity to mitochondrial metabolism (Groten & MacVicar, 2022; Stoler et al., 2022), allowing neurons to rapidly adapt to changing energy demands. Previously, we found a striking enrichment of MCU in hippocampal area CA2 (Pannoni et al., 2023; Farris et al., 2019), a subregion of the hippocampus that is critical for social memory (Hitti and Siegelbaum 2014; Stevenson and Caldwell 2014; Dudek, Alexander, and Farris 2016). In contrast to CA1 neurons, CA2 neurons are known for being resistant to tetanus-induced LTP (Zhao et al., 2007), likely due to robust calcium buffering and extrusion mechanisms (Simons et al., 2011; Carstens and Dudek, 2019). While this is the case for CA3 synapses onto the proximal dendrites of CA2 (Schaffer collaterals; SC), entorhinal cortex layer II (ECII) synapses onto distal dendrites of CA2 (perforant path; PP) readily express LTP (Chevaleyre and Siegelbaum 2010; Dudek et al. 2016). There are not any known candidates distinguishing CA2 proximal and distal synapses that could mediate these functional differences in plasticity. We recently uncovered that the propensity for LTP in CA2 correlates with more mitochondrial mass and higher expression of MCU selectively in distal dendrites compared to proximal dendrites (Pannoni et al., 2023). The distinct enrichment of MCU in CA2 distal dendrites is not solely due to more mitochondrial mass, as other mitochondrial markers, such as COX4, do not show the same level of enrichment as MCU in CA2 distal dendrites and do not differ between areas CA1 and CA2 (Pannoni et al,. 2023). Thus, we hypothesized that layer-specific enrichment of MCU in CA2 distal dendrites may be important for promoting plasticity in otherwise LTP-resistant CA2 neurons. Here, we generated a conditional KO of MCU in CA2 neurons and found MCU is necessary for LTP at ECII-CA2 distal synapses, but the lack of LTP at CA3-CA2 proximal synapses is unaffected. MCU deletion caused mitochondrial fragmentation across all CA2 dendritic layers and therefore layer-specific mitochondrial structural diversity was unaffected. These data expand on the underexplored role of MCU at the post-synapse in a cell type critical for social memory. Ultimately, understanding how diverse mitochondria regulate cellular functions to meet cell-type and circuit-specific needs is critical to our overall understanding of mitochondria in brain health and disease.

## RESULTS

### Conditional deletion of MCU in hippocampal CA2 neurons

To examine whether MCU plays a role in the layer-specific plasticity profile of CA2, we generated a CA2- specific conditional knockout mouse of MCU by crossing an Amigo2-cre mouse line (Alexander et al., 2018) to an MCU ^fl/fl^ line (Kwong et al., 2015). Sections from adult MCU ^fl/fl^;Amigo2-cre negative (CTL) and MCU^fl/fl^;Amigo2-cre positive (cKO) mouse brains were immunostained for MCU and CA2 neuronal marker RGS14 to validate the selective loss of MCU in CA2 neurons. On average, 88% (± 2.2, N = 7 mice) of RGS14 positive CA2 neurons express MCU in CTL mice. After postnatal MCU deletion, 10% (± 2.7, N = 8 mice) of RGS14 positive CA2 neurons express MCU (Fig. 1AB). Consistently, there was a significant decrease in the number of RGS14 positive neurons per section that express MCU in cKO mice compared to CTL mice (Fig. 1B; CTL: 39.8 ± 2.6 neurons, cKO: 4.3 ± 1.1 neurons, two-tailed unpaired t-test, p<0.0001) without a difference in the total number of RGS14 positive CA2 neurons per section between genotypes (Fig. 1C; CTL: 45.4 ± 2.6 neurons, cKO: 39.9 ± 3.1, two-tailed unpaired t-test, p= 0.204) MCU fluorescence intensity was reduced selectively in CA2 neuron cell bodies (Fig. 1D; two-way ANOVA, overall effect of genotype F (1, 52) = 22.45, p<0.0001, sidak’s post hoc test CTL v. cKO CA2 p< 0.0001), with no significant change in MCU fluorescence intensity in cell bodies in CA1, dentate gyrus (DG), or the neighboring cortex in MCU cKO compared to CTL (sidak’s post hoc CTL v. cKO CA1, DG, CTX p> 0.05). Further, a decrease in MCU fluorescence intensity was seen across all layers of CA2 dendrites in the cKO compared to CTL (Fig. 1E). In the neuropil layer, MCU labeling is predominantly detected within dendrites, where it localizes to the inner mitochondrial membrane as visualized with protein-retention expansion microscopy (Fig. 1F).

**Figure 1:**
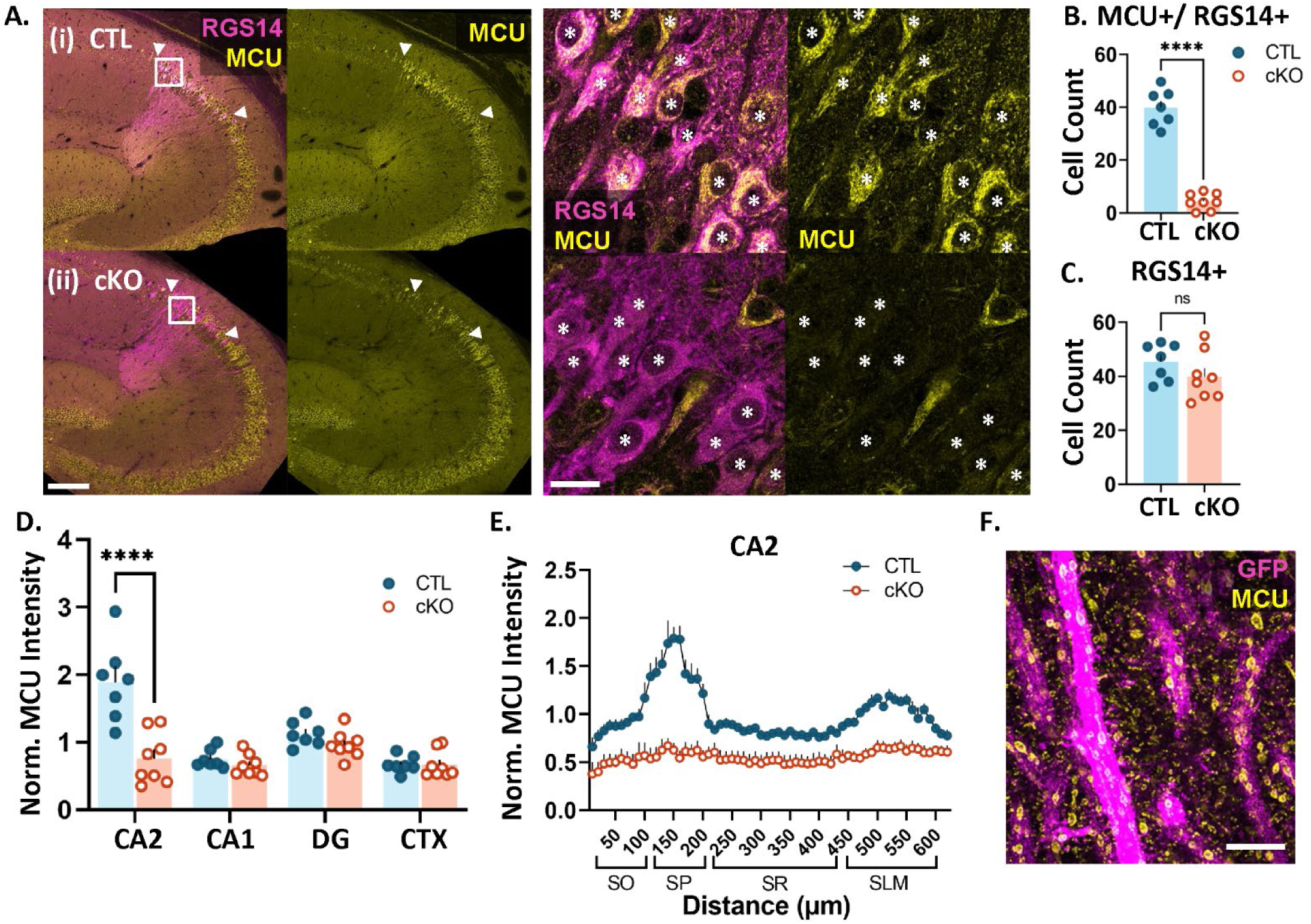
MCU expression is significantly reduced in CA2 neurons of cKO mice. A. Representative images of RGS14 (magenta) and MCU (yellow) immunostaining in CTL (i) and cKO (ii) mice. *Right:* Higher magnification images of CA2 neurons. Asterisks indicate RGS14-positive CA2 neurons. B. Quantification of the number of RGS14+ CA2 neurons per section expressing MCU (CTL: 39.8 ± 2.6 neurons, N=7 mice, cKO: 4.3 ± 1.1 neurons, N=8 mice, two-tailed unpaired t-test, p<0.0001) C. The total number of RGS14-positive CA2 neurons per section does not differ between CTL and cKO mice (CTL: 45.4 ± 2.6 neurons, cKO: 39.9 ± 3.1, two-tailed unpaired t-test, p= 0.204) D. MCU fluorescence intensity in CTL and cKO mice in CA2, CA1, dentate gyrus (DG) and neighboring cortex (CTX). Data were normalized to the CTL average. (two-way ANOVA, overall effect of genotype F (1, 52) = 22.45, p<0.0001; overall effect of subregion F (3, 52) = 17.80, p<0.0001, interaction F (3, 52) = 12.74, p<0.0001; sidak’s post hoc test CTL v. cKO CA2 p<0.0001). E. Line plot analysis of MCU fluorescence intensity across CA2 layers in CTL and cKO mice. Data are normalized to the CA2 CTL average. Representative super resolution image of MCU (yellow) in genetically labeled CA2 dendrites (magenta) using expansion microscopy (ExM). Scale = 200 µm and 25 µm (A) and 5 µm, ExM adjusted (F). ****p<0.0001

### CA2-specific MCU cKO results in impaired LTP at distal dendrite synapses

Next, to assess the role of MCU in the propensity of CA2 distal synapses to express LTP, we recorded extracellular field potentials (FP) from CA2 neurons in acute hippocampal slices from adult CTL and cKO mice. A stimulating electrode was placed in either the Schaffer collateral (SC) inputs to CA2 stratum radiatum (SR, proximal dendrites) or the perforant path (PP) inputs to CA2 stratum lacunosum moleculare (SLM, distal dendrites) and the recording electrode was placed in the CA2 stratum pyramidal (Fig. 2A). The recording site was marked by ejection of CFDA (a green fluorescent dye) for confirmation of placement within CA2. First, we examined whether baseline synaptic field responses were altered by MCU cKO by comparing stimulus-response relationships (input/output curve) for evoked FP peak amplitudes recorded in slices from CTL and cKO mice. No significant differences were found between CTL and cKO slices in the overall input-output curves for stimulation of either SC inputs to SR (p > 0.05 for all stimulus intensities, two-tailed Welch’s *t*-tests, Fig. 2B) or PP inputs to SLM (p > 0.05 for all stimulus intensities, two-tailed Welch’s *t*-tests; Fig. 2C). Further, there was no significant difference between CTL and cKO slices in the half-maximal FP peak amplitude evoked during baseline recordings (SC inputs to SR: CTL = 200 ± 55 μV, cKO = 165 ± 61 μV, p = 0.61; PP inputs to SLM: CTL = 63 ± 12 μV, cKO = 65 ± 11 μV, p = 0.94; two-tailed Welch’s *t*-tests), or in the stimulation intensity required to produce half-maximal responses (SC inputs to SR: CTL = 25 ± 4 μA, cKO = 22 ± 3 μA, p = 0.55; PP inputs to SLM: CTL = 19 ± 3 μA, cKO = 22 ± 3 μA, p = 0.56; two-tailed Welch’s *t*-tests). These results show that basal synaptic transmission is not altered by MCU deletion in CA2. For LTP induction experiments, a prerequisite stable 10-minute pre-conditioning baseline was obtained at 0.1Hz. This was followed by delivery of a strong tetanizing stimulation consisting of three bursts of high frequency stimulation (3 x 100 Hz for 1 second) with an interburst interval of 10 minutes. Subsequently, post-conditioning responses were recorded for a period of 60 minutes at 0.1Hz. Only one recording was made from each slice. Figure 2D shows the average normalized field potential peak amplitude evoked by stimulation of SC inputs to SR over time in the CTL and cKO conditions. The ratio of the normalized field potential peak amplitude during the last 5 minutes of post-conditioning / last 5 minutes of pre-conditioning (post/pre ratio) is also plotted in Fig 2E. Consistent with observations in wildtype mice (Zhao et al., 2007), stimulation of SC inputs to the SR of CTL mice did not induce a net LTP and this did not change in cKO mice (Fig. 2DE; average post/pre ratio: CTL = 1.07 +/- 0.08, n= 7 slices from 7 mice, cKO = 1.06 +/- 0.07, n= 9 slices from 9 mice; p = 0.45, one-tailed Welch’s *t*-test). In contrast to CTL mice, where stimulation of PP inputs to SLM induced robust LTP as previously described (Chevaleyre and Siegelbaum, 2010), stimulation of PP inputs to SLM failed to induce a robust LTP in cKO mice (Fig. 2FG, average post/pre ratio: CTL= 1.54 +/- 0.19, n= 8 slices from 7 mice, cKO = 1.08 +/- 0.08 n= 7 slices from 7 mice; p = 0.03, one-tailed Welch’s *t-*test). Individual time plots of normalized FP peak amplitude over time are presented for each slice in Supplemental Figure 1. We saw a heterogeneous response to strong tetanizing stimulation with a variety of post/pre ratios (plasticity outcomes). For stimulation of SC inputs to SR in CTL and cKO slices, the majority had responses showing no change (post/pre ratios within +/-10% of baseline), while a single slice in each group exhibited long term depression (LTD, post/pre ratio <90% of baseline), and 33 - 43% displayed weak LTP (post/pre ratio = 114 - 143% of baseline). This heterogeneity resulted in a net no change from baseline is consistent with previous reports with various stimulation protocols (Zhao et al 2007). In contrast, stimulation of PP inputs to SLM in CTL slices 75% showed robust LTP (post/pre ratios = 120 - 248% of baseline), and none showed LTD; while in cKO slices 57% displayed weak LTP (post/pre ratios = 112 - 138% of baseline) and 14% displayed LTD. These results confirm layer-specific plasticity profiles in CTL CA2 and demonstrate that MCU deletion impairs the capacity of synapses in SLM to undergo robust LTP.

**Figure 2:**
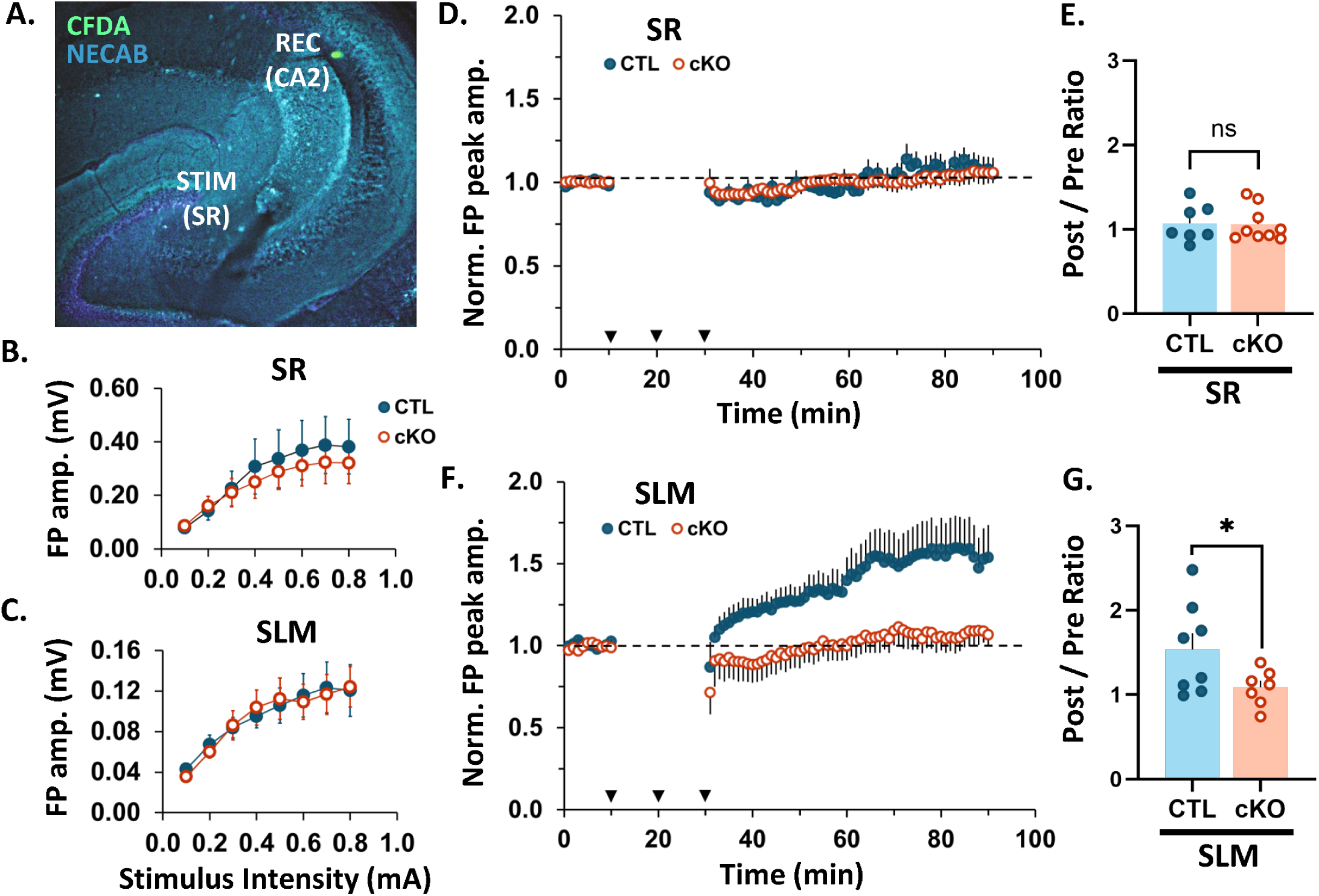
CA2 MCU cKO blocks LTP in CA2 SLM, with no effect in plasticity-resistant CA2 SR. **A.** Representative image of the recording and stimulating site in an acute hippocampal slice. CFDA dye (green) was pressure ejected from the recording pipette. NECAB staining (cyan) delineates CA2. **B.** Input-output curve showing the relationship between field potential amplitude and stimulus intensity after stimulation of SC inputs to SR of CTL (blue, closed) and cKO slices (orange, open). No significant difference in response amplitude at any stimulus intensity between CTL and CKO slices with t-test. N = 7 CTL, 9 cKO slices. **C.** Input-output curve after stimulation of PP inputs to SLM of CTL and cKO slices. No significant difference in response amplitude at any stimulus intensity between CTL and CKO slices with t-test. N = 8 CTL, 7 cKO slices. **D.** Average time plot of normalized field potential peak amplitudes evoked in CA2 by stimulation of SC inputs to SR of CTL and MCU cKO slices. N = 7 CTL, 9 cKO slices. Dotted black line = baseline (all plots). Error bars = SEM (all plots). Arrow heads = stimulation at 100 Hz for 1s; 3 bursts with 10 min interval. **E.** Post/pre ratio of normalized field potential peak amplitudes evoked in CA2 by stimulation of SC inputs to SR of CTL and cKO slices. One-tailed Welch’s t-test; N = 7 CTL, 9 cKO slices. **F.** *(I.)* Average time plot of normalized field potential peak amplitudes evoked in CA2 by stimulation of PP inputs to SLM of CTL and MCU cKO slices. N = 8 CTL, 7 cKO slices. Post/pre ratio of normalized field potential peak amplitudes evoked in CA2 by stimulation of PP inputs to SLM of CTL and cKO slices. One-tailed Welch’s t-test; N = 8 CTL, 7 cKO slices. **ns** = p < 0.05; **\*** p > 0.05.

**Supplemental Figure 1:**
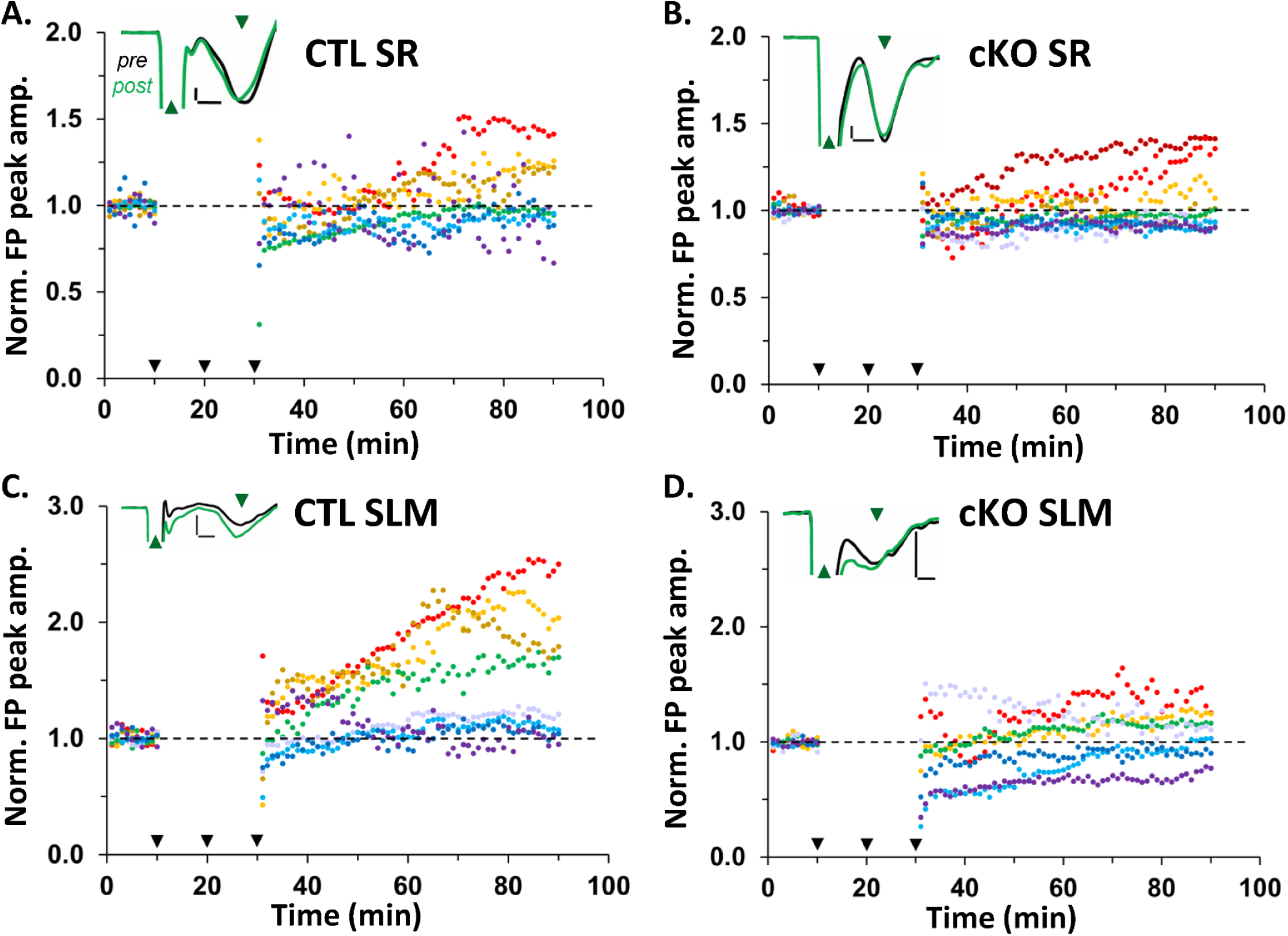
Individual time plots of normalized field potential peak amplitude for each region and genotype showing the heterogeneity of responses by animal. **A.** Individual time plots of normalized FP peak amplitudes evoked by stimulation of SC inputs to SR of CTL slices. Individual plots are color-coded according to post/pre ratio, with progressively warmer colors representing progressively higher post/pre ratios (all panels). Black dotted line = baseline (all panels). Black arrowheads = stimulation at 100 Hz for 1s; 3 bursts with 10 min interval. Inset shows a representative recording of the average evoked response recorded during the last 5 minutes of pre-conditioning (black line) versus the last 5 minutes of post-conditioning (green line; all panels). Green up arrowhead = stimulus artifact; green down arrowhead = FP. Scale bars: 0.1mV, 0.5ms (all panels). **B.** Individual time plots of normalized FP peak amplitude evoked by stimulation of SC inputs to SR of cKO slices. **C.** Individual time plots of normalized FP peak amplitude evoked by stimulation of PP inputs to SLM of CTL slices. Individual time plots of normalized FP peak amplitude evoked by stimulation of PP inputs to SLM of cKO slices.

### A trending reduction in spine density in MCU cKO mice is driven by a decrease in immature spines

LTP involves the strengthening of existing synapses as well as growth and stabilization of new spines and synapses (Hill and Zito, 2013; Engert and Bonhoeffer, 1999; Holtmaat and Svoboda, 2009), thus we hypothesized there may be a loss of spines associated with the LTP deficit in CA2 SLM. To test this, we impregnated hippocampal sections from CTL and MCU cKO mice with Golgi-Cox staining solution (Glaser et al., 1981), a mercury-based sparse cell fill, and quantified the density of dendritic spines in CA2 SLM. Figure 3 shows a Golgi-Cox stained hippocampal section highlighting CA2 (Fig. 3A) and representative dendrite segments from CA2 SLM of CTL (Fig. 3B) and cKO (Fig. 3C). Although not significant, there was a trend towards lower spine density in the MCU cKO compared to CTL (Fig 3D, p = 0.052, Welch’s t-test; N = 8 mice per genotype). Spine density per 100 µm decreased from 97.2 (± 2.1) spines in CTL dendrites to 86.5 (±5.5) in MCU cKO dendrites, representing an 11% average decrease in density. There was significantly more animal variability in spine density for MCU cKO mice than CTL mice (p = 0.022, F = 6.77; F test for variance). In a subset of dendrites, spines were qualitatively classified as immature (filopodia and thin spines) or mature (stubby, mushroom, and branched spines) to assess whether the decrease in spine density was equivalent in immature and mature spines, or whether one group was more impacted (arrows in Fig. 3BC). The decrease in total spine density appears to be driven mostly by a decrease in immature spine density in cKO mice compared to CTL (Fig. 3E; average CTL 91.1 ± 6.0 spines/100 µm, cKO 80.7 ± 4.6 spines/100 µm, p= 0.065, Mann Whitney one-tailed t-test) without changes in mature spine density (Fig 3F; average CTL 31.0 ± 3.32 spines/100 µm, cKO 34.9 ± 3.85 spines/100 µm, p= 0.232, unpaired one-tailed t-test). Due to the trending reduction in overall spine density in the cKO, the relative percentages of immature and mature spine morphologies only differed slightly between cKO and CTL (CTL immature 74.5 ± 2.8%, mature 25.6 ± 2.8%; cKO immature 70.2 ± 2.5%, mature 29.8 ± 2.5%).

**Figure 3:**
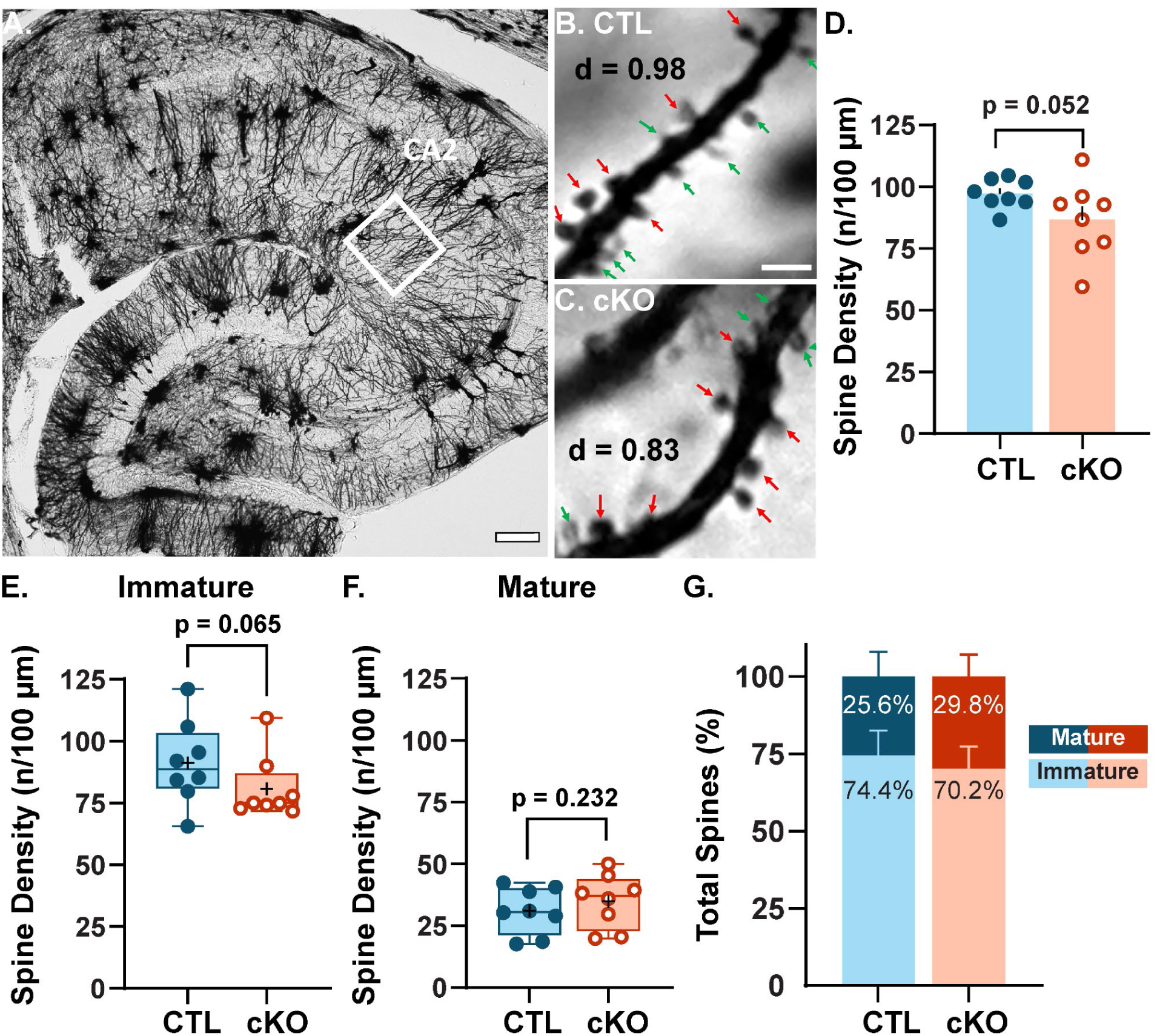
MCU deletion in CA2 decreases the frequency of immature spines and increases the variability of total spine density. **A.** Representative Golgi-Cox stained mouse hippocampus. White box identifies CA2 SLM dendrites. **B.** Representative image of dendritic spines in CA2 SLM from CTL mice. Immature spine = green arrow; mature spine = red arrow; density (d) = number of spine / length of dendrite. **C.** Representative image of dendritic spines in CA2 SLM from MCU cKO mice. **D.** Average spine density in CTL CA2 SLM (blue, closed) and MCU cKO (orange, open) mice. Trending reduction in SLM spine density in MCU cKO compared to CTL (p=0.052, Welch’s t-test; N = 8 mice per genotype). There was a significant increase in variability per animal in MCU cKO mice compared to CTL (p = 0.022, F = 6.77; F test for variance). **E.** Immature spine density per animal in CTL and MCU cKO mice (p= 0.065, Mann Whitney, one-tailed t-test, N=8 mice per genotype). Immature spines consisted of combined counts for filopodia and thin shaped spines. **F.** Mature spine density per animal in CTL and MCU cKO mice (p= 0.232, unpaired one-tailed t-test, N=8 mice per genotype). Mature spines combined counts for stubby, mushroom, and branched shaped spines. Distribution of mature and immature spine morphologies in CTL and MCU cKO mice represented as percent of the total spine count. Scales = 100 µm (A) and 2 µm (C). + denotes average.

### CA2 MCU cKO alters the morphology and mitochondrial content of dendritic mitochondria

Studies have shown the importance of mitochondria both pre- and post-synaptically to support synaptic function and plasticity (Li et al., 2004, Stowers et al., 2002; Smith et al., 2016; Rangaraju et al., 2019). We previously showed that the propensity for LTP at CA2 distal dendrite synapses corresponds with a layer-specific enrichment of MCU and larger mitochondria relative to the other dendritic layers (Pannoni et al., 2023), which in theory could produce more ATP (Lewis et al., 2018). Calcium influx into mitochondria regulates both mitochondrial bioenergetics and the balance of fission / fusion (Llorente-Folch et al., 2015; Zhao et al., 2015), thus we hypothesized that MCU deletion may lead to changes in mitochondrial ultrastructure that could explain a plasticity deficit in CA2.

To look at changes in mitochondrial morphology after MCU deletion, we compared mitochondrial ultrastructure in scanning electron microscopy images from CTL and cKO mice. We acquired images from large (150 x 150 μm^2^) regions of interest (ROIs) from CA2 stratum oriens (SO), SR and SLM of each genotype. A deep learning AI was used to selectively segment dendritic mitochondria in a subset of image tiles from each ROI (Biodock, 2023). A total area of 155,200 µm^2^ was analyzed with the AI over 2 days. The AI correctly identified an estimated 92.2% of dendritic mitochondria, with an error rate of 2 errors / 100 µm^2^ area.

In CTL mice, we noted mitochondrial structural heterogeneity across CA2 dendritic layers (Supplemental Fig. 2A). Mitochondria in SO of CTL mice were uniquely rounded relative to SR and SLM, with an aspect ratio closer to 1 (Supplemental Fig. 2B; Two-way ANOVA with non-parametric post hoc; Aspect Ratio SO vs SR: p < 0.0001; SO vs SLM: p < 0.0001; N SO = 2353, SR = 2904, SLM = 5236 mitochondria). We confirmed our previous findings based on manual segmentation that mitochondria in SLM of CTL CA2 are significantly larger in area than mitochondria in SR (Supplemental Fig. 2C; Area SR vs SLM: p < 0.0001) (see Pannoni et al., 2023). In CA2 SLM, the distance between neighboring mitochondria was significantly less than mitochondria in SO or SR (supplemental Fig 2D; NN Distance SR vs SLM: p < 0.0001; SO vs SLM: p < 0.0001). Mitochondria were also longer in CA2 SLM (supplemental Fig 2E; Feret’s Diameter SR vs SLM: p < 0.0001; SO vs SLM: p < 0.0001), and there was greater overall mitochondrial content, as measured by mitochondrial count and total mitochondrial area per 100 μm^2^ (Supplemental Fig. 2F, Count SR vs SLM: p < 0.0001; SO vs SLM: p < 0.0001; Total Area SR vs SLM: p < 0.0001; SO vs SLM: p < 0.0001; N CTL SO = 223, SR = 279, SLM = 260 100 µm^2^ area tiles). Thus, mitochondria in the proximal (SR) and distal (SLM) dendrites are separated by their mass, diameter and overall content (Supplemental Fig. 2F), whereas mitochondria in the basal dendrites (SO) are separated mostly by their aspect ratio (Supplemental Fig. 2B).

**Supplemental Figure 2:**
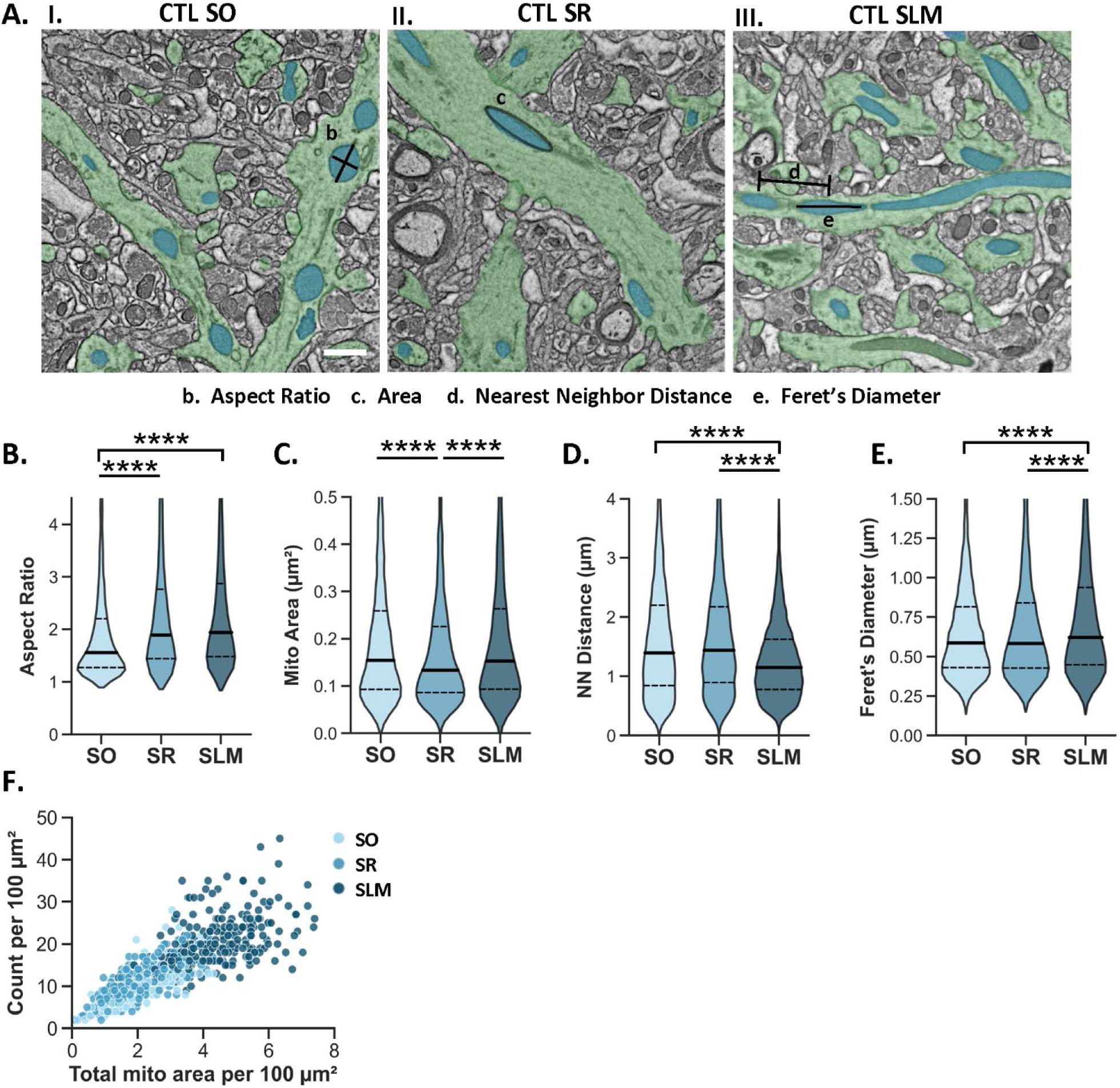
Morphometrics separate mitochondria into distinct populations within CA2 dendritic layers **A.** Representative SEM images showing dendritic mitochondria (blue) and dendrites (green) in CA2 SO, SR and SLM of CTL mice. Examples of the measured metrics are illustrated. Scale bar = 1 μm. **B.** Mitochondrial aspect ratio in CA2 SO, SR, and SLM of CTL. Overall effect of layer was significant (two-way ANOVA, F (2, 23653) = 203.2, p < 0.0001; SO n = 2353, SR n = 2904, SLM n = 5236 mitochondria, 3 mice). **C.** Individual mitochondria area in the same dataset as in (B). Overall effect of layer was significant (two-way ANOVA, F (2, 23653) = 131.7, p < 0.0001). **D.** Mitochondria nearest neighbor distance in the same dataset as in (B). Overall effect of layer was significant (Two-way ANOVA, F (2, 23653) = 570.00, p < 0.0001). **E.** Mitochondria Feret’s diameter in the same dataset as in (B). Overall effect of layer was significant (two-way ANOVA, F (2, 23653) = 162.6, p < 0.0001). **F.** A correlation of mitochondrial count and total mitochondrial area per 100 μm^2^ image tile in CA2 SO, SR, and SLM. Overall effect of layer was significant for both mitochondria count (two-way ANOVA; F (2, 1551) = 590.35, p < 0.0001) and total mitochondria area (two-way ANOVA; F (2, 1551) = 976.34, p < 0.0001; SO n = 223, SR n = 279, SLM n = 260, n = 100 μm^2^ tiles). Mann Whitney post hoc tests and Sidak’s correction comparing layers are shown. For all violin plots, solid line = median; dashed line = upper and lower quartiles.

Due to the hierarchical nature of this dataset, assuming that individual mitochondria are statistically independent may result in false positives. To address this, we applied Bayesian statistics with a hierarchical bootstrap (Saravanan et al, 2020) (Supplemental Fig. 3). The data were randomly resampled (with replacement) at the level of animal, hippocampal section, image tile and individual mitochondria, and the median of the resampled data was calculated for each layer in CA2. This was repeated 10,000 times to get a population of resampled medians for each group, which were plotted together on joint probability distribution plots for each comparison. Consistently, the median mitochondria size and mitochondria count per tile were greatest in CA2 SLM of CTL compared to SR (Supplemental Fig. 3CD, Area: 83% of iterations; Supplemental Fig. 3GH, Count: 100% of iterations). Collectively, these data confirm that the AI detects the established morphological diversity of mitochondria across the dendritic layers of CA2.Mann Whitney post hoc tests comparing genotypes are shown on the plots.

**Supplemental Figure 3:**
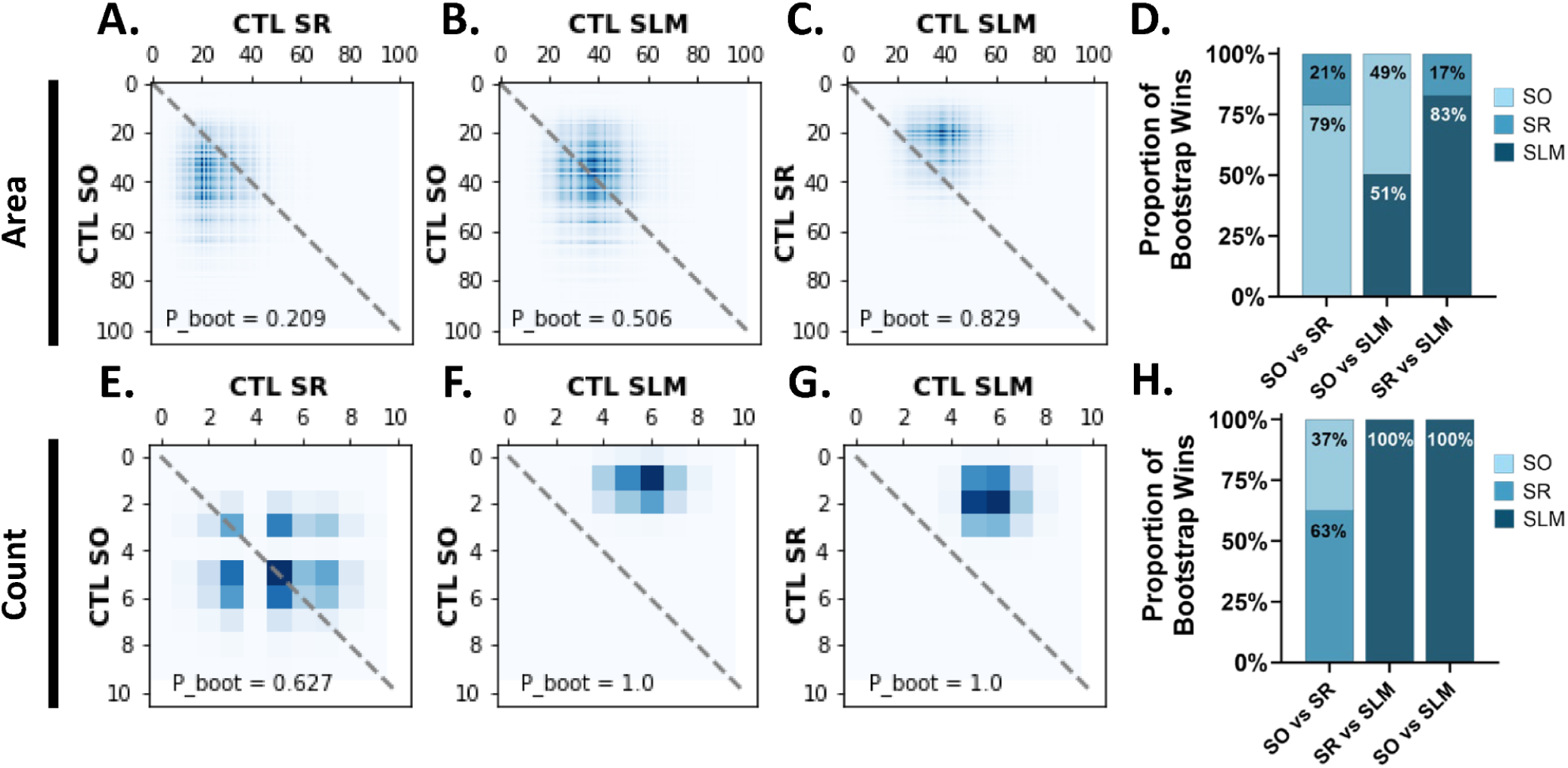
Confirmation of mitochondrial ultrastructure differences across dendritic layers of CTL CA2 with a hierarchical bootstrap. **A-C.** Comparison of resampled mitochondria area medians across SO and SR (**A**), SO and SLM (**B**) or SR and SLM (**C**) of CTL CA2 with a probability distribution plot. P_boot = the proportion of medians within the upper triangle of the plot. N = 10,000 repetitions. Data was binned into 100 equal bins. **D.** Bar plot showing the proportion of bootstrap repetitions where mitochondria area was greater in SO, SR or SLM for each comparison. **E-G.** Comparison of resampled median mitochondria count per tile across SO and SR (**E**), SO and SLM (**F**) or SR and SLM (**G**) of CTL CA2. N = 10,000 repetitions. Data was binned into 10 equal bins. **H.** Bar plot showing the proportion of bootstrap repetitions where mitochondria count per tile was greater in SO, SR or SLM for each layer comparison in CTL CA2.

To determine whether loss of MCU affects mitochondrial ultrastructure, we compared dendritic mitochondria morphology in each layer of CA2 in CTL and MCU cKO mice (Fig. 4). In MCU cKO mice, individual mitochondria were significantly smaller (Fig. 4D; two-way ANOVA with sidak’s post hoc test; effect of genotype p < 0.0001; N CTL SO = 2353, SR = 2904, SLM = 5236 mitochondria, cKO SO = 2860, SR = 4007, SLM = 6299 mitochondria) and shorter in the long axis (Feret’s diameter, Fig 4E; effect of genotype p < 0.0001) across all dendritic layers compared to CTL mice, with no changes in aspect ratio (Fig. 4F, effect of genotype p = 0.566). This indicates that mitochondria are smaller in both dimensions in cKO relative to CTL. Unexpectedly, there was a decrease in nearest neighbor distance (Fig. 4G; two-way ANOVA with sidak’s post hoc test; effect of genotype p < 0.0001; N CTL SO = 223, SR = 279, SLM = 260, cKO SO = 223, SR = 279, SLM = 260) and an increase in mitochondrial count per 100 μm^2^ in all dendritic layers of cKO mice (Fig. 4H; effect of genotype p < 0.0001). A decrease in individual mitochondria area in combination with an increase in mitochondria count suggests that MCU cKO causes an increase in mitochondria fragmentation. The increase in mitochondrial count was greatest in SLM (Median number of mitochondria per 100 μm^2^: CTL SLM: 19 ± 0.24, cKO SLM: 23 ± 0.27; CTL SR: 10 ± 0.41, cKO SR: 13 ± 0.34; CTL SO: 10 ± 0.30, cKO SO: 13 ± 0.36; Fig 4H) and appears to be driving an increase in overall mitochondrial content after MCU cKO (Fig. 4H).

**Figure 4:**
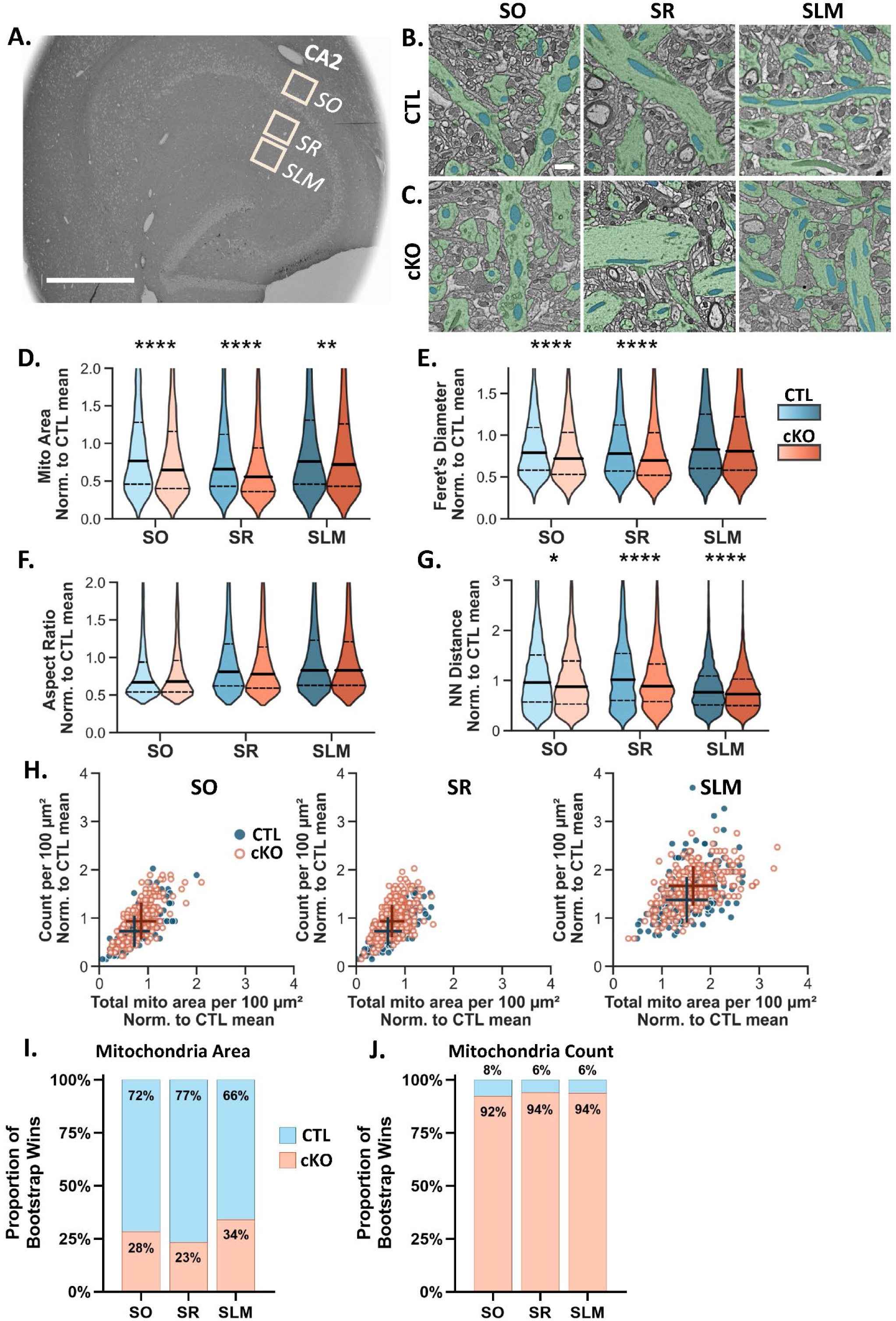
Dendritic mitochondria in MCU cKO mice are smaller and more numerous than CTL mice. **A.** Representative scanning electron microscopy (SEM) overview of a horizontal hippocampal section. Boxes represent the sampling areas from CA2 SO, SR and SLM. Scale = 600 µm. **B.** Representative SEM images from CA2 SO, SR, and SLM of CTL showing dendritic mitochondria (blue) and dendrites (green). Scale = 1 µm for B-C. **C.** Representative SEM images from CA2 SO, SR and SLM of the MCU cKO. **D.** Violin plots of individual mitochondrial area in 100 μm^2^ image tiles from CA2 SO, SR and SLM in CTL (blue) and cKO mice (orange). All plots were normalized to the overall mean of the CTL. Two-way ANOVA, significant overall effect of genotype F (1, 23653) = 49.06, p < 0.0001; significant overall effect of layer F (2, 23653) = 131.67, p < 0.0001; interaction F (2, 7) = 7.04, p < 0.001; CTL SO n = 2353, CTL SR n = 2904, CTL SLM n = 5236. cKO SO n = 2860, cKO SR = 4007, cKO SLM = 6299 mitochondria. Solid line = median; dashed line = upper and lower quartiles. **E.** Normalized mitochondria Feret’s diameter from the same dataset in (D). Two-way ANOVA, significant overall effect of genotype F (1, 23653) = 25.47, p < 0.0001; significant overall effect of layer F (2, 23653) = 162.55, p < 0.0001; interaction F (2, 23653) = 4.15, p < 0.05. **F.** Normalized mitochondria aspect ratio from the same dataset in (D). Two-way ANOVA, significant overall effect of layer F (2, 23653) = 203.19, p < 0.0001 but no overall effect of genotype F (1, 23653) = 0.566, p > 0.05. See Supplemental Fig 2 for CTL layer pairwise comparisons. **G.** Normalized mitochondria nearest neighbor distance from the same dataset in (D). Two-way ANOVA, significant overall effect of genotype F (1, 23653) = 109.63, p < 0.0001; significant overall effect of layer F (2, 23653) = 570.00, p < 0.0001; interaction F (2, 23653) = 14.83, p < 0.0001) **H.** Correlation plot comparing normalized mitochondrial count and normalized total mitochondrial area per 100 μm^2^ in CA2 SO, SR, and SLM in CTL (blue, closed) and MCU cKO (orange, open). Overall effect of genotype was significant for count (two-way ANOVA, overall effect of genotype F (1, 1551) = 105.47, p < 0.0001; significant overall effect of layer F (2, 1551) = 590.35, p < 0.0001, CTL SO n = 223, CTL SR n = 279, CTL SLM n = 260. cKO SO n = 218, cKO SR = 298, cKO SLM = 279, n = 100 μm^2^ tiles) and total area (two-way ANOVA, overall effect of genotype F (1, 1551) = 27.06, p < 0.0001; significant overall effect of layer F (2, 1551) = 976.34, p < 0.0001). + indicates the median of each group with horizontal and vertical lines indicating standard deviation for each axis. **I.** The proportion of bootstrap iterations (n = 10,000) where median mitochondria area was greater in the CTL (blue) or the cKO (orange) for each layer of CA2. The proportion of bootstrap iterations (n = 10,000) where the median number of mitochondria per tile was greater in the CTL (blue) or the cKO (orange) for each layer of CA2.

We also performed the hierarchical bootstrap analysis to compare mitochondria area and count per tile in CTL and cKO mice (Figure 4IJ; Supplemental Fig. 4). Joint probability distributions of the bootstrap medians were plotted comparing across layers in the cKO (Supplemental Fig. 4A-H) and comparing CTL and cKO in each layer (Supplemental Fig. 4I-N). The resampled median mitochondria area was smaller in the cKO than the CTL in SO in 72% of the bootstrap repetitions, 77% for SR, and 66% for SLM (Fig. 4I; Supp. Fig. 4I-K), indicating that there is a higher probability that mitochondria are uniformly smaller in the cKO compared to CTL. The resampled median number of mitochondria per tile was greater in the cKO than the CTL in SO in 92% of the bootstrap iterations, 94% for SR, and 94% for SLM (Fig. 4J; Supp. Fig. 4L-N), indicating that there is a very high probability that mitochondria are consistently more numerous in the cKO compared to CTL.

**Supplemental Figure 4:**
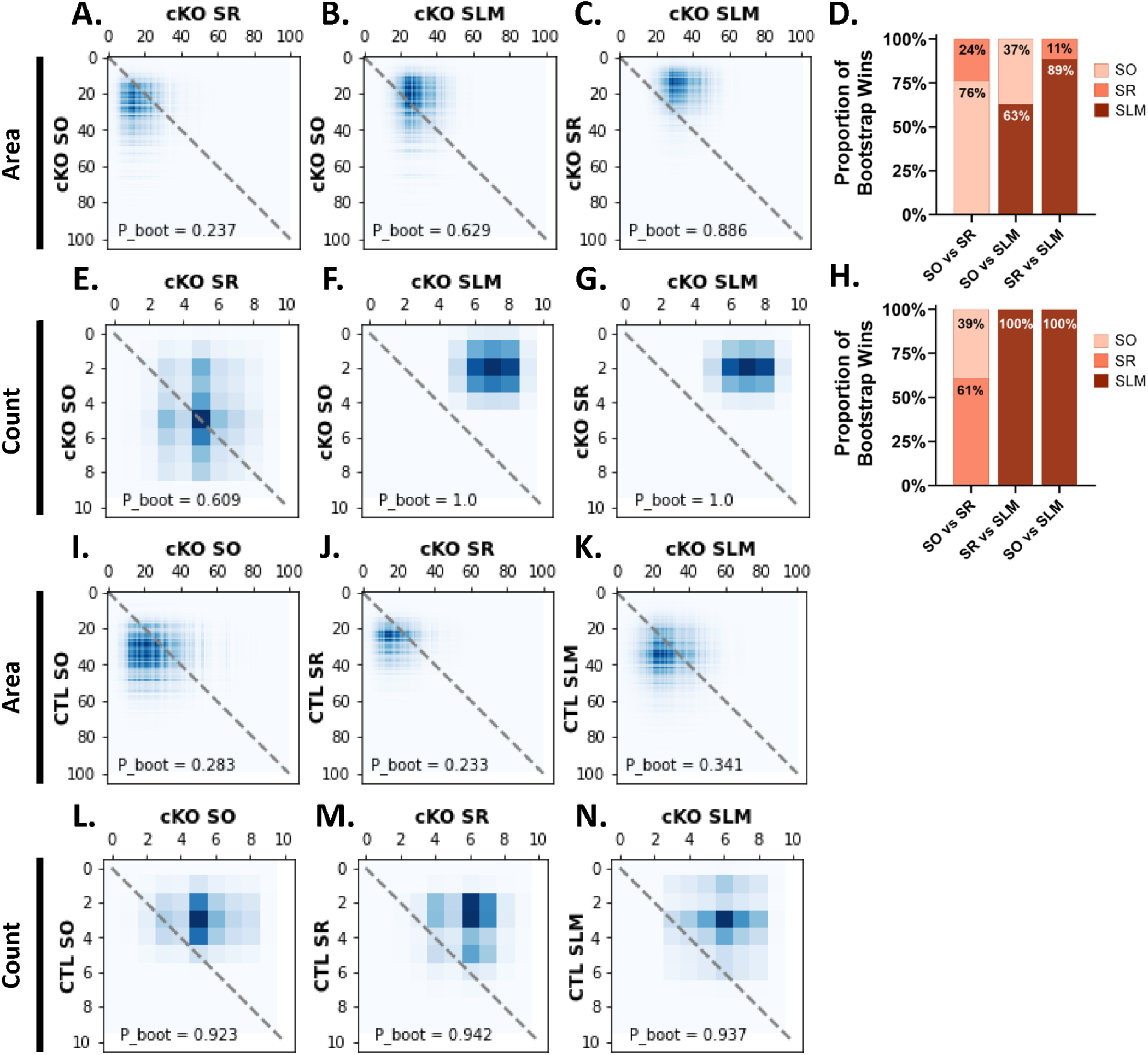
Confirmation of mitochondrial ultrastructure differences across dendritic layers of CTL and cKO mice with a hierarchical bootstrap. **A-C.** Comparison of resampled mitochondria area medians across SO and SR (**A**), SO and SLM (**B**) or SR and SLM (**C**) of CA2 in MCU cKO mice. P_boot = the proportion of medians within the upper triangle of the plot. N = 10,000 repetitions. Data was binned into 100 equal bins. **D.** Bar plot showing the proportion of bootstrap repetitions where mitochondria area was greater in SO, SR or SLM for each layer comparison in the cKO. **E-G.** Comparison of resampled median mitochondria count per tile across SO and SR (**E**), SO and SR (**F**) or SR and SLM (**G**) in CA2 of MCU cKO mice. N = 10,000 repetitions. Data was binned into 10 equal bins. **H.** Bar plot showing the proportion of bootstrap repetitions where mitochondria count per tile was greater in SO, SR or SLM for each layer comparison. **I-K.** Comparison of resampled mitochondria area medians in the cKO and CTL for SO (**I**), SR (**J**) and SLM (**K**). This is the same bootstrap population as in Figure 4I. **L-N.** Comparison of resampled mitochondria count medians in the cKO and CTL for SO (**L**), SR (**M**) and SLM (**N**). This is the same bootstrap population as in Figure 4J.

Taken together, our data suggest that mitochondrial fragmentation is increased in MCU cKO in CA2 compared to CTL, and this effect was seen across all dendritic layers. Strikingly, the relative differences in mitochondria size and overall content across dendritic layers were not altered in MCU cKO mice compared to CTL (CTL: Supplemental Fig. 3DH; cKO: Supplemental Fig. 4DH). Although we saw no obvious signs of damaged or unhealthy mitochondria in the MCU KO condition at the ultrastructural level, there may be functional changes or deficits that would not be observed in EM micrographs, such as decreased metabolism and ATP production. We did note several examples where the distribution of mitochondria within the dendrites changed from a linear network with mitochondria oriented end to end in the control dendrites, to a stacked distribution resembling multiple lanes of traffic in many cKO dendrites (see Fig. 4C), which is supported by a decrease in the nearest neighbor distance (Fig. 4G). It is not yet clear the functional significance, if any, of this change.

## DISCUSSION

In the present study, we examined the role of MCU in LTP, dendritic spine density and mitochondrial ultrastructure in CA2 dendrites using a CA2-specific MCU knockout mouse line. We found that MCU is necessary for LTP at CA2 distal synapses, but MCU loss does not alter the lack of plasticity at CA2 proximal synapses. The LTP deficit at synapses in CA2 distal dendrites of MCU cKO correlated with a trending decrease in overall spine density compared to CTL distal dendrites, which was predominantly accounted for by fewer immature spines. Looking at the effect of MCU cKO on dendritic mitochondria ultrastructure in CA2, mitochondria were smaller, more numerous and closer together in MCU cKO mice, suggesting there is more mitochondrial fragmentation. These ultrastructural changes were seen across all dendritic layers of CA2 to different extents. Accordingly, the loss of MCU did not alter the layer-specific differences in mitochondria morphology across CA2 dendrites, suggesting that the asymmetrical expression of MCU across CA2 dendritic layers is not necessary to establish or maintain the structural diversity of mitochondria. However, MCU enrichment in CA2 SLM may still confer unique functional properties to mitochondria that are necessary for LTP at those synapses. While plasticity at the ECII-CA2 synapse has not yet been functionally linked to social memory, activity at the lateral ECII-CA2 synapse is essential for social memory (Lopez-Rojaz et al., 2022). Recent evidence links mitochondrial function to social behavior and deficits (Ülgen et al., 2023), however whether MCU-enriched mitochondria in CA2 contribute to its role in social memory remains to be tested.

### A role for MCU in the propensity of CA2 distal synapses to undergo LTP

It was recently shown that MCU is required for action potential evoked production of NADH by mitochondria in acute cortical slices (Groten and MacVicar, 2022). While these data focus on the role of MCU in powering sustained action potential firing, others have shown mitochondrial calcium uptake in cortical neuron dendrites from acute slices selectively occurs with the coincidence of pre- and postsynaptic activity, suggesting that dendritic mitochondria may respond selectively to plasticity-inducing stimuli (Stoler et al., 2022). Thus, it is possible that MCU plays a general role in proper LTP expression by boosting ATP, but differential MCU expression may tune the mitochondrial response to the unique calcium dynamics of the cell or circuit. Indeed, high-frequency action potential firing causes MCU-dependent mitochondrial calcium uptake that differs between the hippocampus and cortex (Groten and MacVicar, 2022), suggesting that variability in the coupling of activity and mitochondrial calcium uptake due to differences in MCU expression could be a mechanism for scaling energy production to meet cell-type specific needs.

Many features of CA2 neurons are unique compared to neighboring hippocampal neurons. For example, CA2 neurons highly express a number of genes that act as molecular brakes on plasticity (Carstens and Dudek 2019), and some of those mechanisms involve restricting calcium signaling (Evans et al., 2018; Simons et al., 2011). LTP is highly dependent on calcium influx into the post-synapse and resulting downstream signaling cascades (Nicoll, 2017). CA2 neurons have a significantly faster rate of calcium extrusion than CA1 and CA3 neurons, as well as an increased calcium buffering capacity, which mitochondria may contribute to (Simons et al., 2009). The same study also showed that CA2 neurons have the lowest endogenous free calcium at rest and that increasing intracellular calcium levels permits LTP at the typically resistant CA2 SR synapses. Because none of the identified brakes on CA2 plasticity are spatially restricted to SR, it is possible that SLM harbors additional mechanisms to overcome these molecular brakes and express LTP. MCU could fulfill this role, as MCU expression is enriched in SLM compared to SR and one proposed function of MCU is to couple neuronal activity to energy metabolism by decoding intracellular calcium levels (Groten and MacVicar, 2022, Stoler et al., 2022; Bas-Orth et al., 2020). We speculate that the enrichment of MCU at CA2 distal dendrites may promote LTP at ECII-CA2 synapses by enhancing the sensitivity of mitochondria to changes in cytoplasmic calcium, potentially to couple it to ATP production.

Because the CA3-CA2 synapse in SR is resistant to LTP, the lack of an effect of MCU deletion on synaptic plasticity in SR was unsurprising. However, another important function of MCU is to buffer intracellular calcium and one might expect an impact on CA2 synaptic responses at either synapse. Indeed, we previously found that acute blockade of MCU with Ru360 in the patch pipette leads to LTD at CA3-CA2 SR synapses in response to an LTP pairing protocol that normally elicits no change at these synapses (Farris et al., 2019). At the time, we suggested MCU may contribute to plasticity resistance of synapses in SR by taking up calcium into the mitochondria and thereby limiting cytoplasmic calcium (Farris et al., 2019). However, now we know that MCU is more enriched near ECII-CA2 synapses, which may have differences in calcium dynamics compared to CA3-CA2 synapses in SR, leading to MCU deletion having a different effect across circuits. In addition, acutely blocking calcium influx pharmacologically would cause transient changes in cytoplasmic calcium levels that are likely not recapitulated in a postnatal genetic deletion model. It is also possible that MCU may differentially impact plasticity depending on the cell types and forms of plasticity involved. Consistent with this idea, global MCU haploinsufficiency enhanced presynaptic LTP at DG-CA3 synapses in hippocampal slices (Devine et al., 2022). Specifically, clearance of presynaptic calcium by mitochondria was reduced in MCU+/- mice, increasing vesicle release probability, despite reduced ATP (Devine et al., 2022). This contrasting finding could be explained by differences in the role of cytosolic calcium versus ATP in presynaptic and postsynaptic LTP (but see also Rangaraju et al., 2014), or due to differences in the effects of cell-specific homozygous MCU deletion mice versus global haploinsufficient MCU mice. While emerging evidence supports a role for MCU in regulating energy production and calcium buffering in both pre (Ashrafi et al., 2020) and post (Stoler et al., 2022) synaptic functions, further studies are needed to resolve its impact on specific cell types and circuits as well as the underlying mechanisms linking it to plasticity.

### MCU loss results in changes to spine morphology

We detected a ∼10% decrease in spine density in the distal dendrites of MCU cKO mice that could contribute to the deficit in LTP. This non-significant decrease is largely explained by a loss of immature spines, while mature spines were considerably stable compared to CTL. This suggests that immature spines may be more vulnerable to loss of MCU. It is widely accepted that mushroom spines are relatively more stable than filopodia and immature spine types (Kasai et al., 2003). Notably, we found that ∼75% of spines in CA2 SLM were immature. This is much higher than estimates from others that ∼20% of spines are immature in adult brains (Berry and Nedivi, 2017), highlighting that CA2 SLM has a particularly high proportion of immature spines, or that our criteria for manually categorizing spine shape differs. However, the LTP deficit may also be due to non-structural changes in dendritic spines that would not be captured by our methods. Spines in CA2 SLM receive input from ECII, and activity from the lateral ECII to CA2 is required for social recognition memory (Lopez-Rojas et al. 2021). Given the LTP and spine deficits we uncovered in SLM, further studies are warranted to test whether MCU cKO in CA2 has an effect on social recognition memory.

### Mitochondrial fragmentation due to MCU deletion

Mitochondria are highly dynamic organelles that are shaped by two opposing forces: fission and fusion. Mitochondrial fission, the division of mitochondria to make new or recycle damaged mitochondria, is mediated by the phosphorylation of dynamin-related protein 1 (DRP1; Zheng et al., 2017). Whereas fusion of mitochondria is mediated by optic atrophy 1 (OPA1) and mitofusins (Legros et al., 2002, Cipolat et al., 2004). The balance of these forces determines mitochondrial form, which is intimately linked to bioenergetics. For example, it’s been shown that larger axonal mitochondria in cortical neurons can produce more ATP (Lewis et al., 2018). Fragmented mitochondria, which could be due to either less fusion or more fission (Detmer and Chan, 2004; Chen et al., 2003), could result in functional consequences, such as decreased ATP production, altered ROS generation or calcium-induced calcium release, that might underlie a plasticity deficit. There are multiple potential causes of mitochondrial fragmentation after MCU deletion. In cultured rat hippocampal dendrites, mitochondrial fission is rapidly triggered by chemical LTP (Divakaruni et al., 2018). This mitochondrial fission requires CaMKII and DRP1 and precedes mitochondrial calcium uptake (Divakaruni et al., 2018). Expression of dominant negative forms of DRP1 blocks mitochondrial fission and LTP in both culture and CA1 of acute hippocampal slices (Divakaruni et al., 2018). Therefore, loss of MCU may elevate cytoplasmic calcium levels and lead to aberrant activation of DRP1 that could, in theory, result in more fragmented mitochondria if fission can occur without mitochondrial calcium uptake. The effect of MCU loss on mitochondrial number was greatest in SLM, suggesting the mitochondria might be more fragmented in SLM relative to SO or SR in MCU cKO mice– although, it did not scale proportionally with a greater decrease in individual mitochondria area or diameter in SLM.

On the other hand, inhibition of MCU has been shown to prevent mitochondrial fission and fragmentation in cultured hippocampal neurons during ischemia (Zhao et al., 2015). MCU has also been shown to mediate mitochondrial fission in rat cortex (Liang et al., 2014). Although these studies are in contrast to our findings that loss of MCU results in an increase in mitochondria fragmentation in CA2, most have looked at MCU loss in the context of injury and disease. Endogenous MCU expression is also highly variable across cell types, thus different cell types may have different sensitivities to MCU-induced mitochondrial fragmentation and cell death (Granatiero et al., 2019). It is also possible that the fragmentation we found is not due to increased fission mediated by DRP1, but instead an impairment in fusion, or a result of unhealthy or damaged mitochondria. Lewis et al showed that a loss of synapses correlated with increased local mitochondrial fission and ULK2-dependent mitophagy in CA1 apical dendrites in an Alzheimer’s disease model (Lewis et al., 2022). However, we did not observe a loss of mitochondrial biomass and the mitochondria in our MCU cKO mice appear ultrastructurally normal in electron micrographs. Even so, the mitochondrial structural changes we observed (i.e. fragmentation) may indicate functional changes that would not necessarily be observed in electron micrographs.

### Altered mitochondrial distribution in dendrites due to MCU deletion

Generally, the distribution of mitochondria strongly correlates with predicted energy usage, which in neurons is highest at the synapse (Harris et al., 2012). In this study and our previous study, we found that CA2 distal dendrites (SLM) harbor more mitochondrial mass than proximal (SR) and basal (SO) dendrites, suggesting that synapses in distal dendrites require more energy. We reasoned that this might be due to the propensity of CA2 SLM synapses to undergo LTP. However, in MCU cKO mice that fail to produce robust LTP at SLM synapses, the relative increased mitochondrial mass in SLM remains. Indeed, the structural heterogeneity across dendritic layers in MCU cKO was similar to CTL, except that mitochondria were overall smaller and more numerous across all dendritic layers in MCU cKO mice (Figure 4 and Supplemental Figure 4). Interestingly, mitochondrial diameter did not decrease in SLM as it did in SO and SR of cKO mice. However, the relative ultrastructural differences across dendritic layers were unchanged. This indicates that MCU expression is not related to the differences in mitochondrial structure across CA2 dendrites. Similar heterogeneity in mitochondrial structure has also been reported in CA1 neurons, with CA1 SLM harboring larger mitochondria than CA1 SR (Virga et al., 2024), suggesting this may be a general pyramidal neuron phenomenon. Virga et al. showed that the difference in mitochondria shape across CA1 dendritic compartments is activity-dependent and regulated by layer-specific differences in AMPK and CAMKK2, which regulate fission/fusion factors. Increasing neuronal activity with picrotoxin induced mitochondrial fission in CA1 dendrites, while acute silencing of CA1 neurons led to an elongation of mitochondria throughout CA1 dendrites. This is in contrast to our findings in CA2, where CA2 SLM has larger mitochondria than SR, despite having a higher synaptic drive and propensity for plasticity relative to CA2 SR (Chevaleyre and Siegelbaum, 2010). This discrepancy could be due to a difference in the balance of fission/fusion factors and/or differences in calcium dynamics between CA1 and CA2 neurons, highlighting that mitochondrial morphology is differentially regulated in different cell-types and circuits. Further studies will be needed to directly compare mitochondrial structure and the expression of fission/fusion factors across CA2 and CA1 dendrites.

We also found that the distances between neighboring mitochondria was reduced in MCU cKO mice across CA2 dendrites, suggesting that the distribution of mitochondria may be altered. The distribution of mitochondria in dendrites is critical for synaptic function (Li et al., 2004). While mitochondria are rarely seen inside dendritic spines in mature neurons, spines are seen to contain Endoplasmic Reticulum (ER) associated with nearby mitochondria in electron micrographs from CA1, CA3, and dentate gyrus (DG) of ground squirrels (Popov et al., 2005). Mitochondria are known to form close connections with ER (< 200 nm) at zones called “mitochondrial associated membranes” (MAM), which allow for communication and the transfer of proteins, ions and metabolites between mitochondria and ER (Rizzuto et al., 2012; Giorgi et al., 2019). An estimated 20% of mitochondrial surface is in close apposition to the ER in living HeLa cells (Rizzuto et al., 1998). It is thought that calcium release from the ER at MAMs is taken up into the mitochondria via MCU and voltage-dependent anion channels on the outer mitochondrial membrane (Rizzuto et al., 2012). MCU deletion could potentially alter these MAM domains and disrupt the ER-mitochondria connection, which would likely have functional consequences at the synapse. In vivo studies in CA1 place cells found that disrupting the ER-mitochondria tether in CA1 pyramidal neurons increased intracellular calcium release and altered the integrative properties of their apical dendrites (O’Hare et al., 2022). This could be one possible explanation for the altered distribution of mitochondria we found within CA2 SLM dendrites in MCU cKO mice, including a decrease in nearest neighbor distance and a “stacked” orientation of the mitochondria in some dendrites. The resolution was not high enough in our SEM dataset to segment or quantify ER-Mitochondria contacts. Mitochondria are also seen to form filamentous reticular networks between other mitochondria in dendrites from CA1, CA3 and dentate gyrus (Popov et al., 2005), which could also be disrupted by MCU cKO. While this has not been studied in CA2 of the hippocampus, we observed what appear to be thin connections between dendritic mitochondria at both the electron microscopy and immunohistological level.

### Limitations

We acknowledge that in the present study we have shown correlations, but not direct causal relationships, between changes in mitochondria morphology and LTP or spine changes. It’s possible these effects occur by different, potentially independent, mechanisms. Further studies will be needed to determine whether MCU deletion in CA2 negatively impacts mitochondrial metabolism; however, this is technically difficult to measure in situ in such a small subregion in a layer specific manner. It is also unclear whether our results are CA2-specific or could be generalizable to other brain areas. In addition, because the cre-dependent recombination of MCU occurs sometime between the age of postnatal (p)4-14, there is the potential for compensation to occur. Compensation could involve MCU-independent methods of calcium entry into the mitochondria, or a downregulation of calcium efflux. MCU-independent calcium influx has been observed in astrocytic mitochondria (Huntington & Srinivasan, 2021); however, given that MCU loss or inhibition by Ru360 blocks calcium uptake in mitochondria from heart, liver, and neurons (Pan et al., 2013; Hamilton et al., 2018), this is unlikely to be the case. Compensation could also involve a shift in mitochondrial respiration to rely more on the malate-aspartate shuttle (MAS). Studies show there is a reversible inhibition of MAS by MCU activation, caused by increased calcium levels in the mitochondrial matrix of isolated mitochondria from brain, liver, and heart (Contreras and Satrústegui, 2009; Satrústegui and Bak, 2015), suggesting that a metabolic switch could be made to compensate when mitochondrial calcium levels are low. However, this would not explain a deficit in LTP after MCU deletion.

Another potential limitation is that mitochondria ultrastructure in 2D SEM images may not provide a full picture of the volume and shape of the mitochondria. The mitochondria count reported is not a true count of individual mitochondria, but a count of mitochondrial segments in the image. This could artificially inflate the mitochondria count particularly in CA2 SLM, where mitochondria are longer and there is more dendritic branching, both of which could result in individual mitochondria going in/out of plane more frequently and being counted as multiple mitochondria. However, it is unlikely this would be different across cKO and CTL groups. In addition, the AI did occasionally miss dendritic mitochondria (∼ 1 dendritic mito missed per 100 µm^2^) and, less commonly, segment non-dendritic mitochondria (2.6% of segmentented mitochondria). Importantly, the AI performed similarly across cKO and CTL images (Error rate per 100 µm^2^ : 2 in cKO; 1.7 in CTL), which allows us to confidently compare dendritic mitochondria across cKO and CTL mice. In addition, the AI analysis replicated what has been previously shown in wild-type CA2 with a genetic fluorescent mitochondrial tag as well as with manual segmentation of SEM images (Pannoni et al. 2023). Others have made comparisons of 3D mitochondrial ultrastructure in SEM across nucleus accumbens (NA), CA1, cortex and dorsal cochlear nucleus (DCN) (Delgado et al., 2019). Consistent with these results, we qualitatively observed that dendritic mitochondria were large and filamentous compared to axonal mitochondria in our 2D SEM dataset (Supplemental Fig. 2A). It is important to emphasize that the traditional statistical measurements used in Figure 4 and Supplemental Figure 2 overestimate the significance of detected effects by treating individual mitochondria as statistically independent. Other EM studies similarly analyze mitochondrial ultrastructure at the level of individual mitochondria (Smith et al., 2016; Faitg et al., 2021), however, they typically do not have thousands of mitochondria per sample. To account for this limitation and bolster the statistical analysis, we included a hierarchical statistical bootstrap to interpret the results in terms of Bayesian probabilities (Supplemental Figs. 3 and 4). Generating repeated samples of the data with a bootstrap provides an estimate of the variability and uncertainty in the observed data, which boosts confidence in the robustness of the results.

Combined, our results demonstrate a role for MCU in regulating layer-specific plasticity in CA2 distal dendrites and for maintaining proper spine and mitochondrial morphology and distribution in dendrites, but MCU is dispensable for mitochondrial structural diversity across CA2 dendrites. We speculate that MCU may generally function postsynaptically to decode cytoplasmic calcium signals to boost metabolic output leading to long lasting changes in synaptic efficacy and that differences in postsynaptic MCU expression may reflect a general mechanism to tune ATP production in different calcium contexts.

## METHODS

### Animals

All experiments were performed on adult (8-16 week old unless otherwise noted) male and female mice on a C57BL/6J background. MCU CTL and cKO mice were generated by crossing a CA2-specific Amigo2-cre mouse line (Alexander et al., 2018) to an MCU ^fl/fl^ line (Kwong et al., 2015). Resulting heterozygous mice were crossed to produce MCU^fl/fl^;Amigo2-cre positive and negative mice, which were then bred together to produce the experimental animals. Genotypes were confirmed for the MCU floxed or WT allele and the presence or absence of cre using Transnetyx genotyping service. The Amigo2-cre line has been validated for conditional deletion of knocked in floxed alleles, demonstrating cre recombination occurring between postnatal ages p4 and p14 (McCan et al 2019 PMID: 31745235). Mice were group-housed when possible under a 12:12 light/dark cycle with access to food and water ad libitum. All procedures were approved by the Institutional Animal Care and Use Committee of Virginia Tech.

### Electrophysiology: In vitro brain slice preparation and recording

Experiments were performed on litters at ages 10-20 weeks with the experimenter blind to genotype. Cutting and recording solutions were made as described in Helton et al 2019. Mice were deeply anesthetized with 4% isoflurane and decapitated. The brain was rapidly removed and cooled for 2 min in ice-cold cutting solution containing (in mM): 10 NaCl, 2.5 KCl, 10 D-(+)-glucose, 25 NaHCO3, 1.25 NaH2PO4, 2 sodium pyruvate, 195 sucrose, 7 MgCl2, 0.5 CaCl2, and saturated with 95% O2/5% CO2 with a final pH of 7.4. Horizontal slices of the hippocampus were cut at 300 µm using a vibratome (VT1000S, Lecia) and placed in artificial cerebrospinal fluid (ACSF) containing (in mM): 125 NaCl, 2.5 KCl, 1.25 NaH2PO4, 25 NaHCO3, 20 D-(+)-glucose, 2 Na-pyruvate, 2 MgCl2, 2 CaCl2, and saturated with 95% O2/5% CO2 with a final pH of 7.4. Slices were incubated in ACSF at 33±1°C for 20 min and then at room temperature for > 40 min prior to recording.

For recording, slices were transferred to a submerged recording chamber perfused continuously with 3 ml/min of oxygenated ACSF at 33 ± 1°C. CA2 pyramidal neurons were visualized using a Zeiss microscope (Axio Examiner.D1; Zeiss) equipped with a W Plan-Apochromat 40x water immersion lens configured for DODT gradient contrast (DGC) microscopy. A stimulating electrode (model #30213; FHC inc.) was placed in either the SC to stimulate inputs to the SR, or PP to stimulate inputs to the SLM. For recording, glass micropipettes (O.D. 1.5 mm, I.D. 1.12 mm; A-M Systems) were pulled on a vertical puller (PC-10, Narishige) to make field potential (FP) pipettes (1.2 ± 0.5 MΩ). FP pipettes were filled with ACSF and placed in the stratum pyramidal of CA2. CFDA-SE (an amine-reactive cell-permeable fluorescent green dye; Thermo-Fisher) was added to the pipette solution and pressure ejected after recording was completed to allow post hoc identification of the recording site (only recordings made from CA2 were kept for analysis).

Afferent stimulation consisted of constant current square wave pulses 10 - 100 µA in amplitude (set to 50% of maximal FP response) and 100 µsec in duration. Pre-conditioning baseline recordings of evoked FP peak amplitudes were made at a stimulation frequency of 0.1 Hz for 10 min. This was followed by a 20 minute period of synaptic conditioning, consisting of three stimulus trains of 100Hz for 1 sec, interleaved with two 10 min rest periods without stimulation (Dasgupta et al., 2017; “STET” protocol). Finally, post-conditioning evoked FP peak amplitudes were evaluated at 0.1 Hz for a period of 60 min.

All recordings were made with a MultiClamp 700B amplifier, digitized at 20 kHz with a Digidata 1440A and recorded using Clampex 10.7 software (Axon Instruments, Molecular Devices). Recordings of evoked FP peak amplitudes were measured as the difference between baseline and peak (analyzed with a 1 msec smoothing window). Any recordings with an unstable baseline (linear best fit of all preconditioning FP peak amplitudes had an r^2^ > 0.2) were discarded. Only one recording was made from each slice, so that a single stimulation protocol was applied in each case.

Data was analyzed using Clampfit 10.7 software (Axon Instruments, Molecular Devices). We assessed plasticity of the FP response in each slice by calculating a post/pre ratio (the average FP peak amplitude for the last 5 minutes of post-conditioning divided by the average FP peak amplitude for the last 5 minutes of pre- conditioning). We defined LTD as a significant decrease (>10%) in post/pre ratio, LTP as a significant increase (>10%) in post/pre ratio; or no change (not significantly different, or <10% change) in post/pre ratio (p < 0.05, *t*- test). Data displayed as time plots show values for normalized FP peak amplitudes averaged over 1 minute intervals (i.e. each data point represents the average of 6 data points collected at 0.1 Hz).

Post Hoc immunofluorescence staining was used to validate the recording site in area CA2 (Helton 2019 PMID: 30067288). After recording, slices were post-fixed in 4% paraformaldehyde for 12-48 hours then transferred to 1X PBS. Slices were permeabilized and blocked overnight with 3% Normal Goat Serum (NGS) in 1X PBS-0.3% Triton X-100 (0.3% PBST) before 2-3 day incubation with primary antibodies at 4C for PCP4 (1:250, Invitrogen Cat# PA5-52209, RRID:AB_2645298) or NECAB2 (1:250, Novus Cat# NBP1-84002, RRIS:AB_11028373). Slices were washed in 0.3% PBST several times and then incubated with AlexaFluor goat anti rabbit 633 (1:250, Sigma Cat# SAB4600141) overnight. After washes with 0.3% PBST, slices were stained with DAPI (Sigma Aldrich D9542, 1:10,000 in PBS) incubated for 30 min in 60% TDE prior to imaging cleared slices weighted with harps in a 6 well glass bottom plate (Cellvis P06-1.5H-N) on an Leica Thunder microscope at 20X.

### Immunofluorescence

Mice were anesthetized with 150 mg/kg sodium pentobarbital and transcardially perfused with 15-20ml of ice-cold 4% paraformaldehyde. Brains were dissected and post-fixed for at least 24 hours before sectioning 40 μm thick sections in the horizontal plane on a vibratome (Leica VT1000S). All brain sections immunostained with MCU were washed in 1X PBS-0.1% Triton X-100 (0.1% PBST) before they underwent antigen retrieval by boiling free floating sections for 5 min in 1 ml of nanopure water, followed by a permeabilization step in 0.1% PBST for 15 min at room temperature. All sections were then blocked for 1 hour in 5% Normal Goat Serum (NGS) in 0.1% PBST. Sections were incubated overnight at 4C (18-24 hours) with primary antibodies: rabbit-anti-MCU (1:2000, Novus Cat# NBP2-76948, Lot# H660681004, RRID:AB_2924913) and mouse-anti-RGS14 (1:500, NeuroMab Cat# 75–170, RRID:AB_2877352). Sections were then washed in 0.1% PBST several times and blocked for 30 minutes in 5% NGS in 0.1% PBST. Sections were incubated for 2 hours at room temperature in secondary antibodies (1:500, Invitrogen, AlexaFluors goat anti rabbit 546 Cat# A11035 and goat anti mouse 488 Cat# A11029) followed by several washes in 0.1% PBST and a final wash in 1X PBS.

### cKO Validation

Fig. 1 includes MCU histology data from XX mice aged 8-16 weeks and 2 mice aged 32-62 weeks per genotype. The results from older mice did not significantly differ from younger mice. 20X (0.8 NA) 16-bit images of MCU and RGS14 immunolabeling were acquired on a Leica Thunder epifluorescence microscope (Leica DMi 8) using identical acquisition parameters for CTL and cKO and Lightning computationally clearing. MCU fluorescence was quantified in FIJI (v. 2.1.0/1.53c, NIH) (Shindelin et al., 2012) using max projected images from 3-5 sections per animal. CA2 neurons were identified via RGS14 labeling. A line ROI was drawn along the length of CA2 neurons (line length 650 µm, drawn at the end of the mossy fiber track with the start of the line being in SO and the midpoint of the line at the middle of SR), CA1 neurons (line length 700 µm, starting at SO with the midpoint of the line at the middle of SR), and DG granule neurons (line length 300 µm, starting at the granule cell body layer, with up to 75 µm covering the cell body layer when possible and the rest of the line on the molecular layer). Fluorescence values along the line were obtained using the FIJI Analyze and Plot Profile functions. For the neighboring cortex, a 400 µm x 400 µm ROI was cropped out of the original image and fluorescence intensity was measured using the Measure function in Fiji. Fluorescence background noise was subtracted for all regions using the negative control (no primary antibody) sections. Due to differences in the length of the CA regions along the dorsal-ventral axis, some of the ROI lines yielded zero values from the line being beyond the image border. These values were removed before averaging across sections per animal. The data were then binned by 10 microns length and averaged across sections to obtain one average fluorescence by distance line plot per animal. The data were then normalized to the CTL animals run in the same IHC cohort. In order to compare across subregions, the peak binned value representing the cell body layer per section was averaged across sections per animal, and compared across regions such that every animal is represented in each region. MCU cKO was further validated through quantification of CA2 cell count using the Cell Counter tool in Fiji. An equal z-stack size was used for all images and cell bodies that expressed MCU and/or RGS14 were manually counted in each of these z sections. Experimenters were blind to genotype through the analyses and used RGS14 to guide placement of lines and cell counts. However, due to the obvious deletion of MCU expression in CA2 neurons, experimental bias could not be completely eliminated.

### Protein-retention Expansion Microscopy (ProExM)

40 μm horizontal mouse brain sections were immunostained then expanded with 4X protein expansion microscopy (ProExM) as previously described in Campbell et al. 2021 using an *Amigo2*-EGFP line to selectively label CA2. Briefly, sections were treated as described above with the following modification. All sections were washed in PBS and blocked for at least 1 h in 5% Normal Goat Serum (NGS)/0.3% Triton-100x. Antibodies (MCU, 1:500 and Chicken anti-GFP, 1:500 Abcam ab13970) were diluted in blocking solution and sections were incubated for 72+ hours at room temperature (RT). After several rinses in PBS-T (0.3% Triton- 100x), sections were incubated in secondary antibodies (1:500, Invitrogen AlexaFluors, goat anti chicken 488 Cat# A11039 and goat anti rabbit 546 Cat#A11035) for 48h at RT. Prior to imaging, adjacent unexpanded sections that were run simultaneously with expanded sections were washed in PBS-T and mounted under Vectashield fluorescence media to calculate pre-expansion nuclei diameters.

Sections to be expanded were incubated overnight in Acryloyl-X stock/PBS (1:100, ThermoFisher, A20770) at room temperature in the dark. All solutions were prepared as described by Asano et al. 2018. Following incubation, the slices were washed twice with PBS for 15 minutes each at room temperature. The gelation solution was prepared by adding 384 uL of monomer solution, 8 uL 4-Hydroxy-TEMPO inhibitor (1:200 w/v, Sigma Aldrich, 176141), 8uL TEMED accelerator (10% v/v, Sigma Aldrich, T7024), and lastly 8uL of APS initiator (10% w/v, Sigma Aldrich, 248614) for each section. Sections were incubated in the gelation solution for 30 minutes at 4C in the dark. Gelated sections were placed on gelation chambers constructed from microscope slides with coverslips as spacers. Our gelation chambers produce gels with the thickness of a single No. 1.5 coverslip (∼0.15mm thick). The chambers were filled with gelation solution and allowed to incubate at 37C for 2 hours in a humidified container. Following gelation, the gelation chamber was deconstructed and the gel was carefully removed from the chamber using a coverslip and Proteinase K-free digestion solution. Gels were then placed in digestion solution containing proteinase K (8U/mL, New England BioLabs, P8107S) and digested for 4 hours at room temperature.

Gels were stained with DAPI (Sigma Aldrich, D9542; 1:10,000 in PBS) for 30 minutes at room temperature with shaking. Finally, the gels were washed twice with PBS for at least 20 minutes and either stored in PBS at 4C or fully expanded in npH20 for imaging. Images of CA2 SLM dendrites were acquired using 4X Super Resolution by Optical Pixel Reassignment (SoRa) on an inverted spinning disk confocal microscope (Yokogawa CSU-W1 SoRa/Nikon Ti2-E Microscope) equipped with Hamamatsu Fusion BT cameras, and 20X water (0.95 NA. WD 1.0 mm) or 60X oil (1.49 NA. WD 0.14 mm) immersion lenses.

### Scanning Electron Microscopy

This protocol was adapted from the protocol version 2 published by NCMIR at UCSD (Deerinck et al. 2022). Mice were anesthetized with sodium pentobarbital (euthanasia solution, 150mg/kg) and perfused with ice-cold 0.1M cacodylate buffer pH 7.4 containing 2.5% glutaraldehyde, 2% paraformaldehyde with 2mM calcium chloride for 3 minutes. The brain was removed and fixed overnight at 4C in the same fixative before vibratome sectioning (Leica VT1000S) into 350-micron thick sections in the 0.1M cacodylate buffer pH 7.4 with 2mM calcium chloride. Sections were placed back in fixative for microdissection three days later. Hemisected brain sections were placed on wax paper with a drop of fixative and a 2mm x 2mm hippocampal microdissection was obtained per brain and placed back in fixative for further processing. Tissues were postfixed with 1.5 % potassium ferrocyanide plus 2% osmium tetroxide then en bloc stained with incubations in thiocarbohydrazide solution, osmium tetroxide, uranyl acetate, and Walton’s lead aspartate. Dehydration was performed by an ethanol gradient and finished in propylene oxide. Tissues were embedded in Epon 812. The embedded tissues were sectioned to 120 nm (Leica EM UC7 ultramicrotome), mounted on a silicon wafer, and imaged in a ThermoFisher Aprea Volumescope at 2nm pixel size. Three 150 x 150 µm regions of interest were obtained per section (basal, proximal, distal CA2 dendrites) from MCU cKO and CTL mice. A representative EM hippocampal section with ROIs drawn is shown in Figure 3A.

### Analysis of SEM images with Biodock

Regions of interest (ROIs, 150 x 150 µm) in SO, SR and SLM of CA2 in three 120 nm sections from each of three CTL and cKO mice were analyzed. The larger ROIs were acquired as 14 x 20 100 µm^2^ tiles with 10% overlap for stitching (∼12.2 x 8.2 µm). Every 8th tile was selected for analysis to avoid analyzing overlapping tiles. A similar sampling scheme was used for a handful of images that were acquired as 2 x 2 6,700 µm^2^ tiles with 10% overlap, resulting in the same-sized 100 µm^2^ tiles for analysis. Image sampling and processing were done in batches with a custom Python code. Images were inverted if necessary, and a Gaussian blur with a radius of 2 nm was applied to all EM images for training and analysis. Tiles were excluded from analysis if they had poor image quality, significant artifacts or rips, contained only axonal tracts or cell bodies, or were otherwise unfit to analyze. A total of 1,559 tiles (∼156,000 µm^2^) were analyzed across the 6 mice.

To train the Biodock AI, we separated a subset of data (∼1% of the total dataset) for training, making sure to include training data from each layer and genotype in the analysis, as well as undesired elements such as cell bodies, artifacts and blank spaces. We selected an appropriate tile size in Biodock (4000 x 4000 pixels) and labeled all dendritic mitochondria in the training images as an object type class. To be counted as a dendritic mitochondria, the object must have distinct borders, a solid dark fill, and be located inside a clear dendrite segment not bordered by myelin or containing synaptic vesicles. We excluded fragments of mitochondria at image edges or if the identity or location of the object was ambiguous. We proceeded to train the AI on the dendritic mitochondria object class, allowing augmentations such as image flipping and brightness/contrast adjustments. We assessed the AI’s performance with a manual spot check on a test dataset of about 2% of the final dataset. The average number of mistakes per tile was counted by category. Categories included missing dendritic mitochondria, border errors, object merging, detection of axonal or soma mitochondria, and identification of non-mitochondrial elements like myelin. The overall sensitivity of the AI model in correcting identifying dendritic mitochondria was 92.2%.

The trained AI was then used to analyze the final dataset of 3 CTL and 3 cKO mice. We configured the AI project settings to define the analysis metrics of interest and set a confidence threshold of 0.4. A size threshold was applied to exclude mitochondrial objects smaller than 0.01 µm or larger than 2.1 µm, which reflects the minimum and maximum area of visually confirmed mitochondria objects in the dataset. All pixel measurements were converted to microns for length and area, and the aspect ratio was calculated by dividing the short axis by the long axis of each object. Total mitochondrial count and total mitochondrial area were summed per 100 µm^2^ tile, and a nearest neighbor analysis was performed with the KDTree function in the Python package Scipy (Virtanen et al., 2020) to determine mitochondrial euclidean distances. Data for the plots in Figure 3 were normalized to the combined mean of the control. Standard two-way ANOVAs and non-parametric posthoc tests with Sidak’s correction were run with the Pingouin package (Vallat, 2018) on the individual mitochondria for our metrics of interest.

### Hierarchical statistical bootstrap of mitochondria ultrastructure

A hierarchical bootstrap was performed in Python to compare individual mitochondria area and mitochondria count per image tile across dendritic layers in the cKO and CTL. Our hierarchical bootstrap was modeled after Saravanan et al, 2019. For mitochondria count per tile, the SEM data was grouped by genotype and 3 animals from each genotype were randomly sampled with replacement. From those resampled animals, 3 hippocampal sections were randomly sampled with replacement from each animal. To determine the ideal sampling size for image tiles, the bootstrap was run multiple times to randomly sample either 20 tiles (undersampling), 32 tiles (max number of tiles per group), or 50 tiles (oversampling) from each sampled hippocampal section. For each repetition in the bootstrap (n = 1000 for the sampling tests), the median mitochondria count per tile within each group was calculated. Oversampling the tiles reduced the bootstrap variability across trials; thus, for the final bootstrap, 50 tiles were randomly sampled per hippocampal section (n = 450 tiles per group). In general, oversampling the data with replacement in a bootstrap analysis provides a better approximation of the original population’s distribution.

For the mitochondria area, we went on to randomly sample the individual mitochondria objects from each sampled image tile to get the median of the resampled population of mitochondria. Another systematic sampling test was done to determine the ideal number of mitochondria to sample from each tile. The bootstrap was run with a sampling size of either 5, 10, 20, 50 or 100 mitochondria from each of the 50 resampled image tiles. It was determined that 100 mitochondria was the most appropriate sampling size, so the final bootstrap sampled 100 mitochondria from each of 50 sampled tiles to get the medians for each layer (n = 45,000 mitochondria per group). For more robust results, the final bootstrap was run on mitochondria count and mitochondria area with 10,000 repetitions instead of 1,000. A custom Python code was used to plot the distribution of the resampled medians comparing the bootstrap population across layers and genotypes (Supplemental Figure 3). For each probability distribution plot, the medians were binned in linear space into 10 equal bins (for count) or 100 equal bins (for area). For the comparisons of interest, stacked bar plots were also created showing the proportion of bootstrap repetitions where the median for each group in the comparison was greater than the second group.

### Golgi staining

Mice aged 8-16 weeks were perfused and postfixed for at least 24 hours with 2.5% glutaraldehyde/2% paraformaldehyde. Then, brains were dissected in 5 mm^3^ blocks and stained with FD Rapid GolgiStain™ Kit (CAT# PK401) as described in the product manual. A subset of the mice used for Golgi analysis were also used for SEM and processed as 350 µm thick slices as described above. Thick brain slices were wrapped with uniformly thicker 2 mm brain sections and processed as tissue bundles as described in Harris et al. (PMID: 6997641) for Golgi-Cox impregnation using the same kit. Both 5 mm^3^ blocks and 350 µm slices were cryoprotected in 30% sucrose for 4-5 days until they sank. Then they were blocked in OCT and cryosectioned at 40 µm thick in the horizontal plane using Leica Cryostat (Leica CM1860). CTL (n=8) and cKO (n=8) brains were processed in pairs containing each genotype to avoid any potential batch issues. High resolution 63X (NA 1.4) images of CA2 SLM (up to 25μm total on Z-axis, optical section thickness= 0.21μm) were acquired using brightfield microscopy on a Leica Thunder microscope. Individual dendrites in the plane were cropped from the 63X images using Fiji and randomly chosen for analysis. Fiji plugin “Dendritic spine counter” was used to measure dendrite length, total spine count and spine density (number of spines / length of the dendrite) from the cropped dendrites. In summary, ∼200 dendrites were analyzed for spine density in total for both groups. A subset of dendritic segments (∼30-40 μm in length) from both CTL and cKO mice was quantified for spine morphology using the same plugin in Fiji. Individual spines were manually classified based on visual characteristics and features as filopodia, thin, stubby, mushroom and branched (two heads) on 2D average projection images and validated with the Z-stacks to confirm the geometry of spine shapes (Risher et al., 2014) Protrusions from the dendrites that were long and thin were marked as filopodia, and protrusions with a small round tip at the end were classified as thin spines. Thick protrusions with larger, rounded head and relatively narrower neck were marked as mushrooms, and small protrusions without any visible neck were classified as stubby. As the staining is dark and opaque in this technique, spines right above and beneath the dendrites were not visualized and no attempt was made to account for these spines. In total, ∼800 spines were classified per genotype. For plots (Fig 3E & F), thin and filopodia spines were combined as immature spines and stubby, mushroom, and branched spines were combined as mature spine class.

### Statistical analyses

A custom Python code was written to parse the segmented data from Biodock and get the aspect ratio, count, total area and distance to nearest neighbor for the dendritic mitochondria objects. Statistical analyses were done in python (v3.11) or Prism (Graphpad Prism 10) with an alpha of 0.05 considered significant.

Data are presented as animal averages, unless otherwise indicated in the legend. In addition to traditional statistics, a hierarchical statistical bootstrap was performed on the segmented mitochondria data as described above.

## DECLARATIONS

### Authors’ contributions

Conceptualization, SF; Methodology, CE, KEP, DVG, MLC, MMA, NA, LAC, RT, SF, QSF; Formal Analysis, CE, KEP, DVG, MLC, MMA, NA, LAC, RT, SF, QSF; Investigation, CE, KEP, DVG, MLC, MMA, NA, LAC, RT, SF, QSF; Writing – Original Draft, KEP, SF; Writing – Review & Editing, KEP, DVG, MLC, MMA, NA, RT, SF, QSF; Visualization, KEP, MLC, MMA, RT, SF, QSF; Supervision, SF; Funding Acquisition, SF.

### Data availability

Data is available from the authors upon request.

## Acknowledgements

We thank the Virginia Tech animal care staff for their support. The authors also acknowledge resources and support from the Cellular and Molecular Imaging Core, part of the Fralin Biomedical Research Institute at VTC. This research would not be possible without the use of the Thermo Fisher Aprea 2 Serial Blockface Scanning Electron Microscope (NIH award 1S10OD026838-01A1) and Christie Lacy for acquiring the micrographs used for the data shown in Figure 4 Supplemental Figure 2.

