## Supplementary figures and images for "MCU-enriched dendritic mitochondria regulate plasticity in distinct hippocampal circuits"

### Supplemental Figure 1

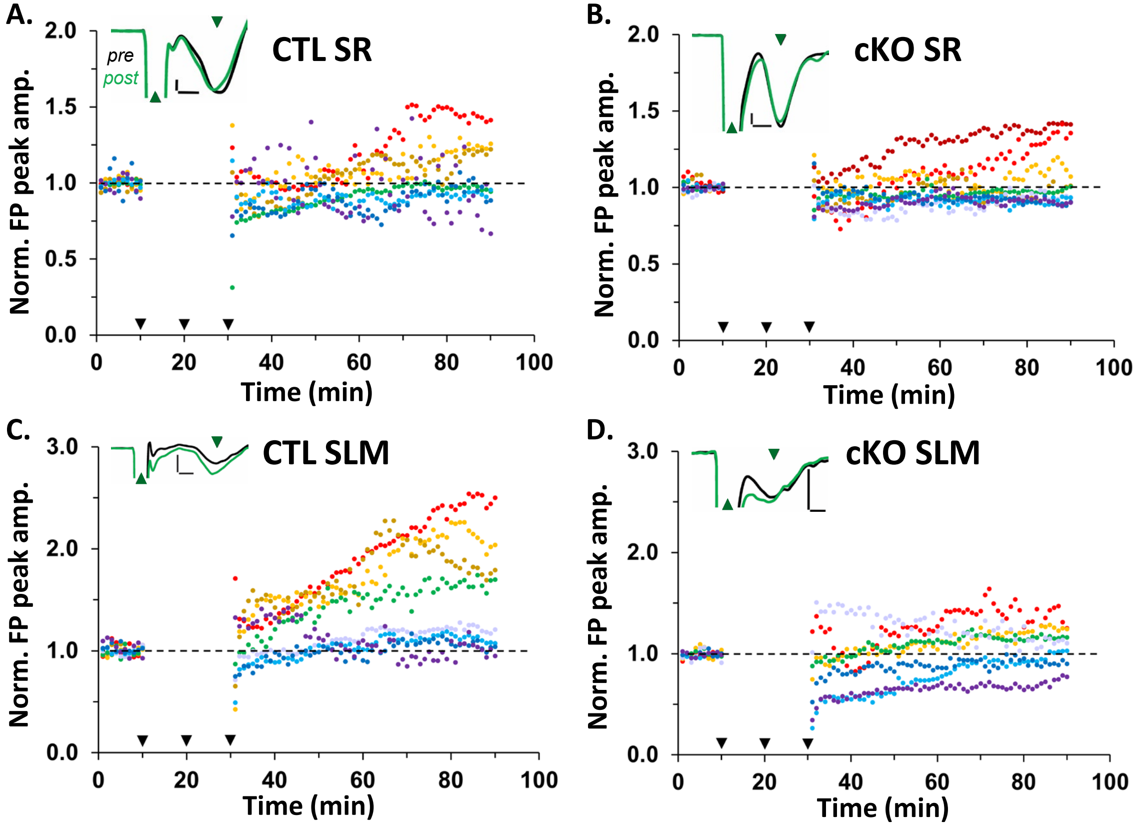

### Supplemental Figure 2

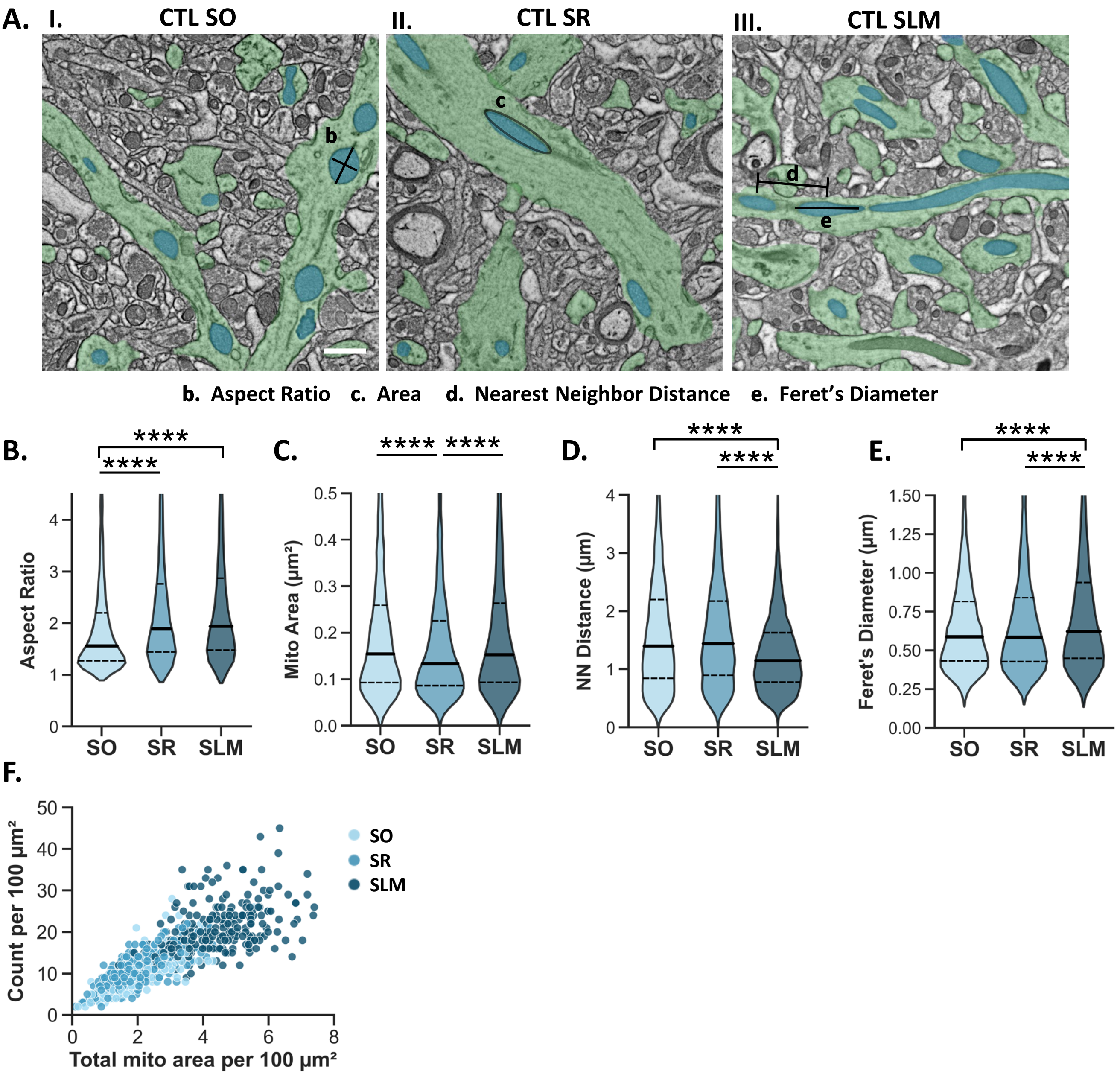

### Supplemental Figure 3

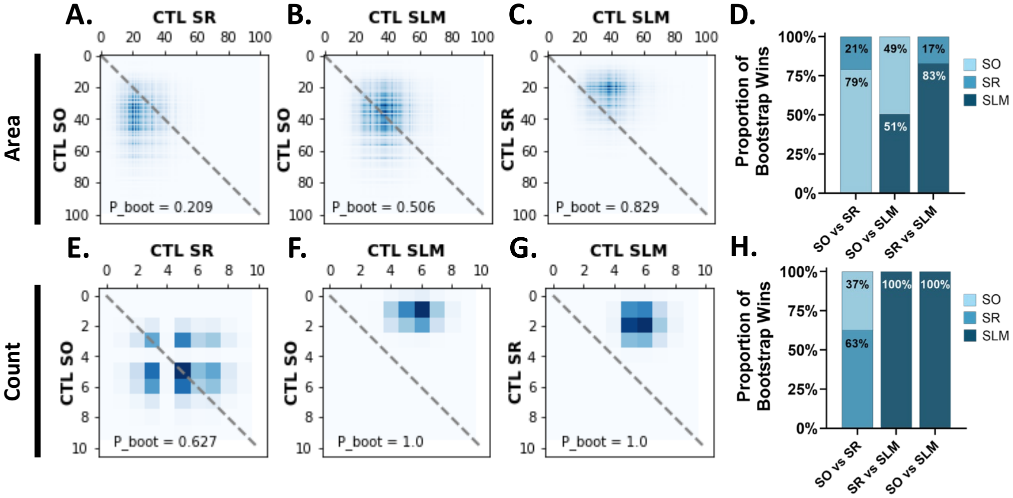

### Supplemental Figure 4

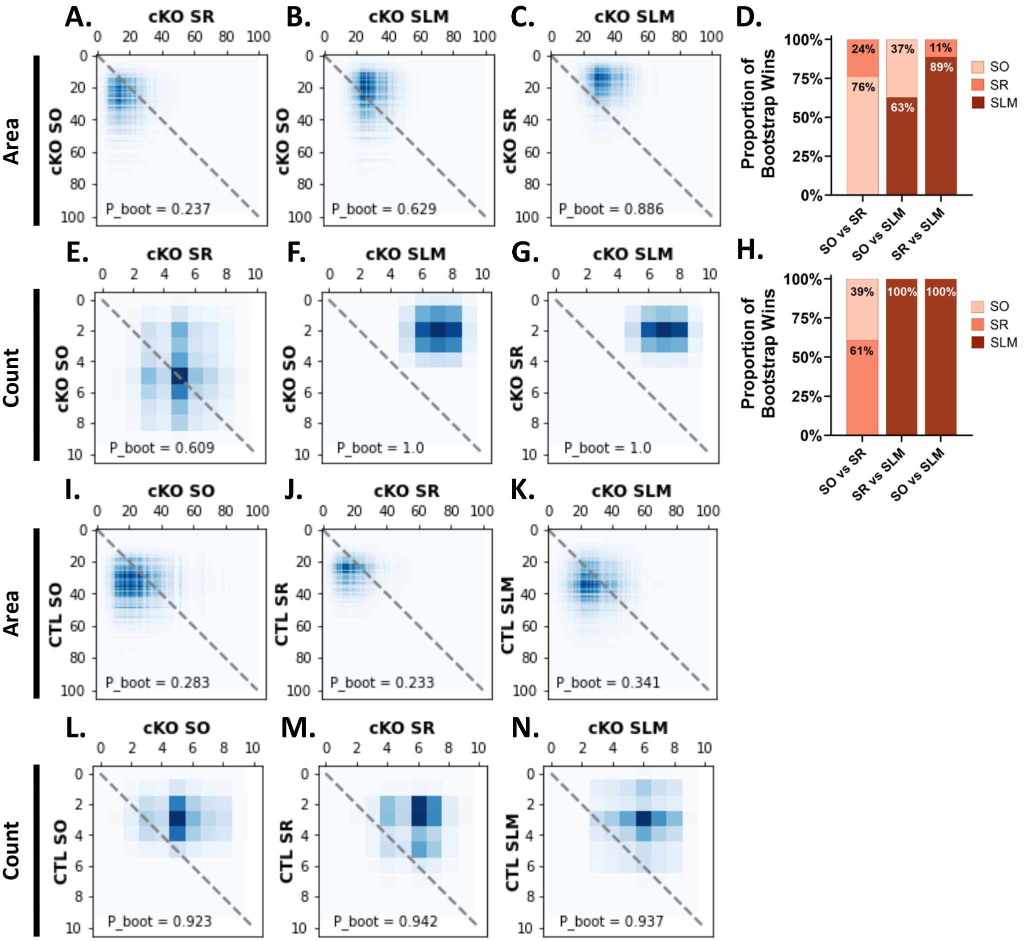
